# Dipteran wing damage discriminates between flight motor inertial-elastic distributions

**DOI:** 10.64898/2026.10.13.757599

**Authors:** Najmeh Asadimoghaddam, Jiaqi Shi, Dorian Decout, Arion Pons

## Abstract

Insects frequently experience irreversible wing damage, yet many maintain flight by adjusting their wingbeat. In Diptera, wingbeat frequency generally increases as wing inertia is lost, but considerably less than predicted by classical models of the flight motor. In this work, we use an inverse-problem approach to identify passive mechanical features of the dipteran flight motor that explain this discrepancy and reproduce experimentally-observed frequency scaling. Combining spatially resolved reconstruction of wing inertia and motor elasticity, finite-element analysis of thoracic deformation, and data-driven viscoelastic models of asynchronous flight muscle, we identify that a compounding interaction between wing root flexion, wing aerodynamics, and muscular viscoelasticity explain the depressed scaling of wingbeat frequency with loss of inertia. Wing inertia is strongly non-uniform and dominated by the vein structure, leading to further depression with respect to loss of area. But, by contrast, thoracic effective inertia is negligible across a wide array of Dipterans and does not contribute to damage-related scaling. Together these results indicate that much of the dipteran flight motor’s responses to wing damage can emerge passively from interactions between the wing, thorax, and muscles; and suggest that a hybrid-elastic architecture with a compliant wing root may be widespread across Diptera. More broadly, we identify wing damage as a tool that can quantitatively discriminate between different candidate models of the insect flight motor, and suggest that altered motor operating conditions can identify mechanical properties difficult to infer from intact flight alone.

## 1 Introduction

Insect wings are delicate exoskeletal structures that cannot heal once damaged—whether by collisions, predator attacks, or gradual wear (Dukas and Dukas, 2011). Such damage can seriously disrupt an insect’s flight: loss of wing area or changes in wing shape can significantly alter the aerodynamic forces generated by the wingbeat (Xie et al., 2026). Across several species, wing damage has been linked to reduced manoeuvrability (Combes et al., 2010; Foster and Cartar, 2011) and hunting and reproductive success (Combes et al., 2010; Vance and Roberts, 2014); increased susceptibility to predators (Jantzen and Eisner, 2008; Vance and Roberts, 2014); and higher overall mortality rates (Allsopp, 1985; Cartar, 1992; Dukas and Dukas, 2011). Nonetheless, several such species also show a remarkable ability to compensate for different forms of wing damage by modifying their wingbeat (Kassner et al., 2016): primarily, increasing wingbeat frequency (Fernández et al., 2012; Sotavalta, 1952), altering wing stroke amplitudes (Kassner et al., 2016; Muijres et al., 2017; Roberts and Cartar, 2015), and changing stroke-plane angle (Kassner et al., 2016).

These compensatory strategies might arise from passive and/or active mechanisms within the insect flight motor. In order Diptera, this motor relies on indirect actuation (Pringle, 1949): the thoracic exoskeleton functions as a dynamic transmission between the primary flight muscles—dorsoventral (DVM) and dorsolongitudinal (DLM)—and the wing. Muscle forces are modulated by thoracic elasticity and damping (Pons, 2025; Pons and Beatus, 2022), and then transmitted through the wing hinge—a specialized joint that functions like a flexible lever—to the wing (Boettiger and Furshpan, 1952). Classically, this flight motor configuration has been considered as a parallel-elastic actuator, in which all thoracic elastic components act in parallel with the muscular actuator, and there is a one-to-one relationship between muscle strain and wing stroke angle (Deora et al., 2017; Lynch et al., 2021; Weis-Fogh, 1973). However, recent evidence increasingly points to series-elastic components in the Dipteran flight motor, manifesting in phase offsets between the muscle and wing (Deora et al., 2021; Dickinson et al., 2005; Fry et al., 2005), and likely arising from flexion at the wing root: such flexion has been directly observed in free flight (Melis et al., 2024).

The interaction between flight motor elasticity—within the thorax and/or the wing root—and muscle activity is an area of both longstanding and recent interest (Gau et al., 2022; Lynch et al., 2021; Machin and Pringle, 1959; Pons, 2023). Dipteran flight primary muscles are asynchronous (Gau et al., 2023): between infrequent neural activations, the muscles continue to activate in response to stretch, generating self-sustained oscillatory contractions when connected to the elastic thorax (Josephson et al., 2000). However, while several mathematical (Gau et al., 2023; Pons, 2023; Wang et al., 2026) and *ex vivo* experimental (Machin and Pringle, 1959; Wold et al., 2023) approaches indicate the significance of thoracic elasticity to wingbeat self-oscillation; and several techniques have studied these elastic properties *ex vivo*, identifying thoracic properties *in vivo* is challenging. In addition to the challenges of small spatial scale and high wingbeat frequency, typically these motors are only observed in one narrow window of operating conditions: tethered, volitional, flight (Chan and Dickinson, 1996; Dickinson et al., 2005; Walker et al., 2014). Within this narrow condition, the types of analyses across oscillation frequency that are typically used for mechanical system identification, *cf*. Lynch et al. (2021) and Wold et al. (2025), are not possible.

We propose that flight motor responses to wing damage can act as an expansion of this window of observation: different states of damage represent mechanically-distinct motor configurations with different wing inertias. Experimental data on wingbeat frequency changes in response to wing wear is reported for a wide range of Dipteran species (Deora et al., 2021; Meng et al., 2023; Muijres et al., 2017; Salem et al., 2022; Sotavalta, 1952), but analysis of this data in the light of flight motor models is limited. Salem et al. (2022) sought to explain trends in wingbeat frequency with respect to wing damage with the predictions of a classical parallel-elastic linear amodel with constant stiffness: under these predictions, wingbeat frequency scales inversely with the square root of inertia. However a comparison between this predicted scaling and wider experimental observations (Sotavalta, 1952) reveals substantial discrepancies (Fig. **1**), with observed scaling substantially depressed with respect to model predictions.

**Fig. 1.**
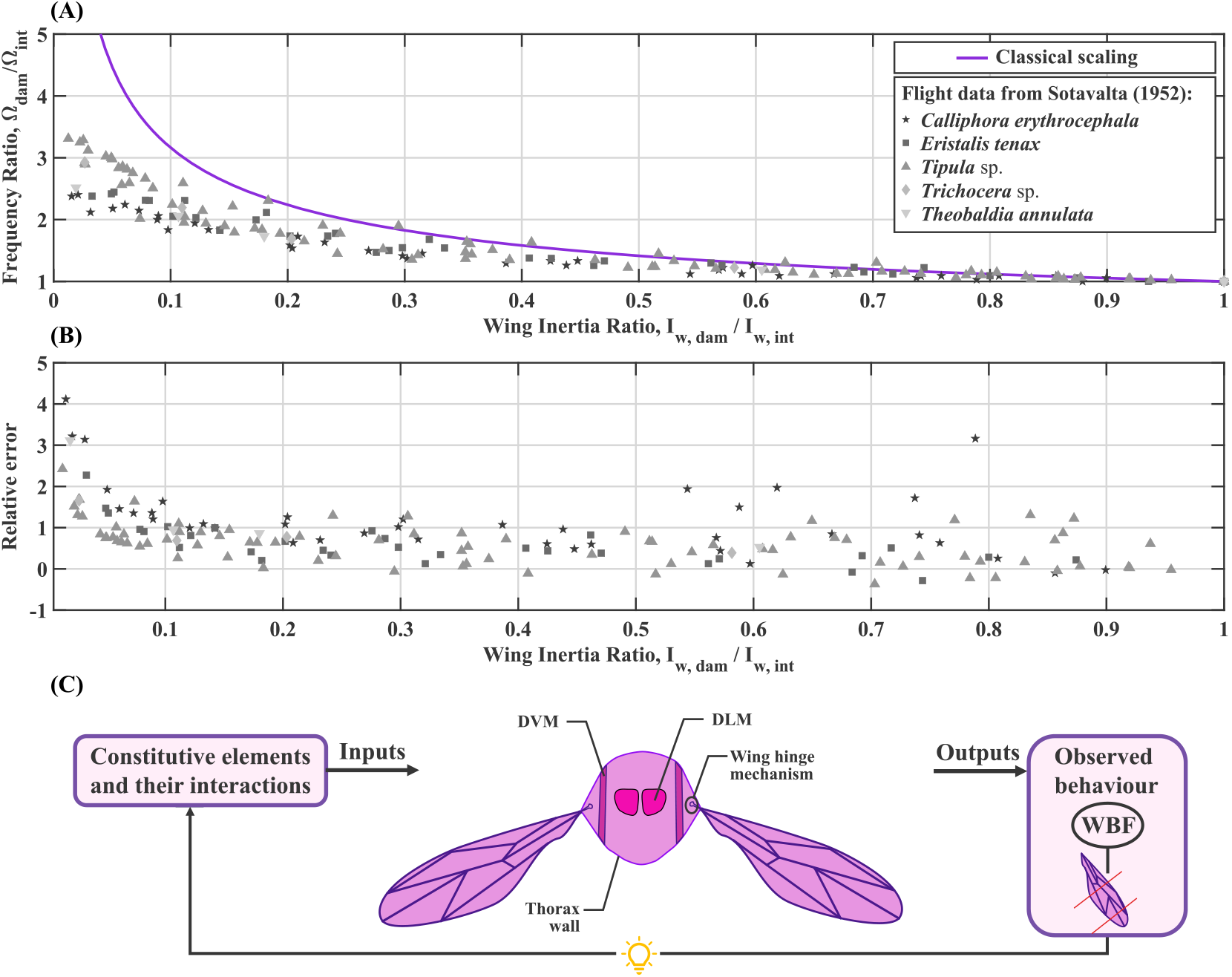
Illustration of depressed wingbeat frequency scaling with wing damage. (**A**) Scaling of wingbeat frequency (Ω) with experimentally-induced wing damage for several dipterans, recorded by Sotavalta (1952): the blowfly *C. erythrocephala*, hoverfly *E. tenax*, a crane fly of *Tipula* sp., a winter crane fly of *Trichocera* sp., and the mosquito *Theobaldia annulata*, now *Culiseta annulata*. Illustrated is the ratio of damaged (Ω_dam_) to intact (Ω_int_) frequency, as a function of the ratio of damaged (I_w, dam_) to intact (I_w, int_) wing inertia, alongside the classical square-root scaling proposed by Salem et al. (2022). (**B**) Factor of error in the classical square-root prediction: positive values indicating that observed scaling is depressed with respect to predicted scaling. Large errors factors are observed—regularly exceeding 1, *i*.*e*., observed data scales half as strongly as predicted. (**C**) Illustration of an inverse-problem approach: wingbeat frequency (WBF) modulations induced by experimentally-induced wing damage (outputs), including the depressed scaling illustrated in (**A**-**B**), are used to infer the underlying constitutive elements of the indirect flight motor (inputs).

In this work, we analyse the contribution of several passive mechanical factors to Dipteran wingbeat scaling under wing damage: the non-uniform distribution of inertia over the wing vein and membrane structure; the inertial contribution of thoracic deformation; the viscoelasticity of the primary flight muscles; and flexion at the wing root. Following an inverse-problem approach, *cf*. Pons et al. (2023), and synthesizing reported X-ray microtomography (μCT) and wing interference pattern (WIP) data with finite-element (FE) simulations, we identify the level to which these factors explain depressed wingbeat frequency scaling. We find that (**i**) thoracic inertia is likely nonsignificant across Diptera; (**ii**) wing inertia is strongly non-uniform and acts to depress wingbeat frequency scaling; (**iii**) individually, muscular viscoelasticity and wing root flexion have a minor contribution to depressed wingbeat frequency scaling, but (**iv**) an interaction between muscular viscoelasticity and wing root flexion leads to an effect that is greater than the sum of its parts, accounting for much of the depressed scaling previously observed. In the light of these results, we predict that wing root flexion—previously, only directly observed in drosophilids (Melis et al., 2024)—is likely present across Diptera; and we offer perspectives on the mechanics behind non-flight vibrations of the flight motor, *e*.*g*., defence buzzing (Vallejo-Marín and Vallejo, 2021). We suggest that further analyses of insect flight motors across different *in vivo* operating conditions have the potential to reveal information that cannot be inferred from only hovering flight.

## 2 Materials and methods

### 2.1 Pixel-wise reconstruction of nonuniform wing inertia

#### 2.1.1 Pixel-wise framework

To quantify the changes in wing rotational inertia effected by wing damage, we develop a method for mapping the distribution of mass and rotational inertia over the Dipteran wing, and apply this method to reported data for *Drosophila melanogaster*. We follow a pixel-wise framework, in which we integrate standard optical imagery, wing interference pattern (WIP) imagery, and X-ray microtomography (μCT) data to build up an estimate of the inertia associated with each pixel of an optical image of the wing. We begin with a high-resolution isotropic RGB image of a *D. melanogaster* wing, obtained online (Jones, 2013). Within this image, we classify pixels as membrane, vein, or image background via combined intensity- and colour-based segmentation: edge detection applied to a contrast-enhanced grayscale image version identifies wing boundaries and internal vein features; colour information from the RGB image is extracted by pixel-wise *k*-means clustering and used to partition pixels into vein, membrane, and background regions. These complementary sources of information are integrated, and refined via morphological operations, to generate a consensus binary wing mask and a processed image with vein, membrane, or background pixels labelled. A schematic overview of the workflow is shown in Fig. **2**A, with further details in the Supplementary Material (§S1.1).

**Fig. 2:**
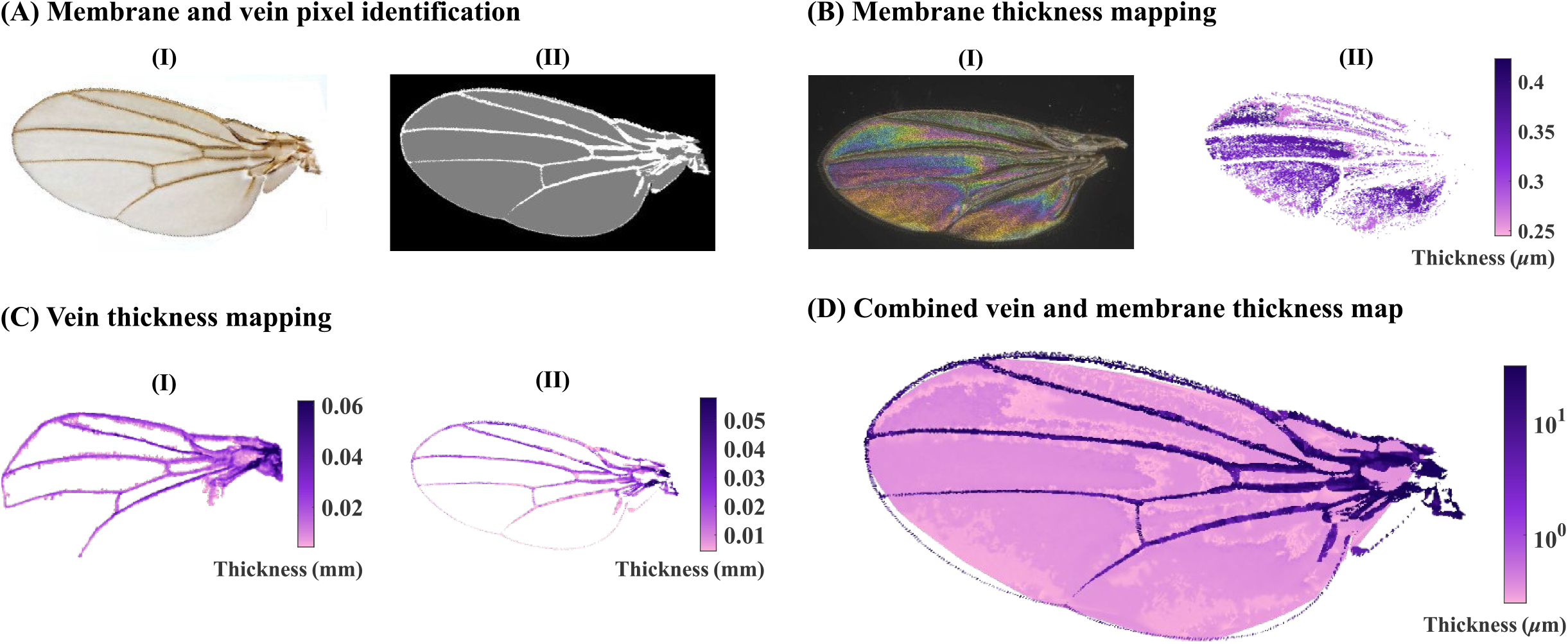
Workflow for pixel-wise reconstruction of nonuniform wing inertia. (**A**) Membrane and vein pixel identification. (A.I) Processed RGB image of a *D. melanogaster* wing used as the input for segmentation (Jones, 2013). (A.II) Labelled image obtained after image processing and refinement, identifying vein, membrane, and background. (**B**) Membrane thickness mapping from WIP imagery. (B.I) WIP of a *D. melanogaster* wing from De Lisle et al. (2025). (B.II) Sparse membrane thickness map obtained from thin-film interference modelling of reconstructed reflectance spectra from the filtered WIP image. (**C**) Vein thickness mapping from μCT data. (C.I) Raw projected vein thickness obtained from the μCT volume. (C.II) Interpolated vein thickness map registered to the original wing image. (**D**) Complete spatially resolved thickness map for membrane and veins.

We then calculate the contributions of the labelled membrane and vein components to wing mass and rotational inertia, over all pixels i, as:

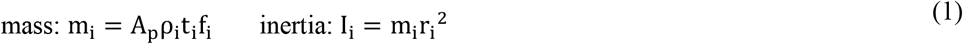

where: t_i_ represents the local thickness of the vein or membrane pixel; A_p_ the physical surface area associated with a single pixel—constant across the image, for an image with isotropic scaling; f_i_ represents the fraction of each pixel’s area that lies within the wing region under consideration—accounting for partial pixel occupancy when pixels located along the wing boundary are not fully contained within the analysed domain; ρ_i_ represents the local volumetric density, differing between vein and membrane; and r_i_ = S_F_(x_root_ − x_i_) is the distance of each pixel centre from the wing root—x_i_ denoting the horizontal position of the pixel centre in the optical image, x_root_ the corresponding position of the wing root, and S_F_ the spatial scale factor converting pixel units into physical units. To perform this computation, we require maps of the wing and vein thickness and estimates of membrane and vein density: to these we now turn.

#### 2.1.2 Membrane and vein thickness mapping

Following Li et al. (2022) we estimate the distribution of membrane thickness over a representative *D. melanogaster* wing via an inverse colourimetry approach applied to wing interference pattern (WIP) imagery recorded by De Lisle et al. (2025) (Fig. **2**B). Invalid (*i*.*e*., unclearly coloured) pixels are excluded using HSV-based and median filtering, and the resulting valid pixels are spatially registered to the anatomical reference image using manually selected control points and a third-order polynomial transformation. For each valid pixel, the recorded WIP RGB triplet is converted to the underlying reflectance spectrum by modelling the camera’s spectral sensitivities as Gaussian functions and estimating the most likely reflectance intensity at each wavelength over the visible wavelength range of 450–650 nm using non-negative least-squares optimization (Davis, 2018). Using constrained non-linear least-squares _ptimization, we fit this reconstructed reflectance spectrum at each pixel to a thin-film interference model describing the transparent membrane reflectance (Li et al., 2022):

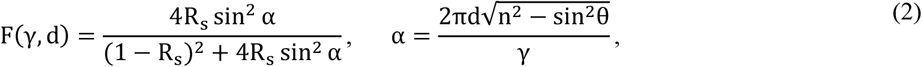

where n = 1.57 is the refractive index of chitin; d is the membrane thickness; θ is the incidence angle, which, for the experimental conditions of De Lisle et al. (2025), is normal (θ = 0°); γ is the wavelength; and R_s_ is the reflection coefficient for s-polarised light:

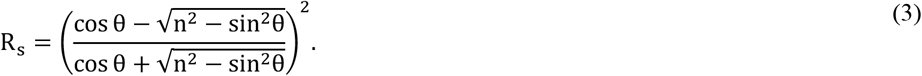

Pixels with poor spectral fits are excluded, and missing values are interpolated using cubic interpolation to preserve the smooth thickness gradients characteristic of thin-film membranes. The resulting thickness field is restricted to the wing domain using the latter’s binary mask. Details of this approach are provided in the Supplementary Material (§S1.2).

To map wing vein thickness, defined as the cross-sectional height of the vein above the membrane, we process *D. melanogaster* μCT data captured by Maisano (2024). We segment the vein structure in 3D Slicer using intensity-based thresholding, producing a two-dimensional thickness map (Fig. **2**C). We resample and register this map to the labelled anatomical wing image using manually selected landmarks and an affine transformation. Nearest-neighbour interpolation is applied to avoid artificial smoothing and preserves local thickness magnitudes along vein structures. Small gaps and undefined pixels, occurring primarily along the boundaries of the valid thickness region, are addressed using a local four-connected-neighbourhood-based filling procedure to ensure spatial continuity along the vein network. The registered thickness map is assigned exclusively to vein pixels (as per §2.1.1) with remaining gaps filled using nearest-neighbour propagation. The final physical vein thickness map overlaid on the original optical wing image is illustrated in Fig. **2**C. Finally, the membrane thickness map and the vein thickness map are combined within the common anatomical reference frame to produce a unified spatial thickness distribution across the wing surface (Fig. **2**D).

#### 2.1.3 Characterization of the Mass Density of the Membrane and Veins

Computing wing mass and inertia from the wing thickness map requires the volumetric densities of membrane and vein structures. We assume the membrane to be composed of a chitin–protein composite with uniform properties, and take its density as 1.2 g cm^−3^, consistent with values for hydrated insect cuticle (Klocke and Schmitz, 2011). However, by contrast, the veins are composite structures consisting of a cuticular shell enclosing fluid- or air-filled internal passageways (Pass, 2018; Sugiyama et al., 2024). Their effective density is thus likely to differ from that of solid cuticle.

We calibrate an appropriate effective vein density by matching model-predicted wing masses to experimental wing mass measurements for *D. melanogaster* (Bergou et al., 2010; Fry et al., 2005; Greenewalt, 1962; Lehmann and Dickinson, 1997; Pétavy et al., 1997). For a range of candidate vein densities, we compute total wing mass as a function of wingspan length using isometrically scaled reconstructed geometry and compare this function to reported trends in mass (Supplementary Material, Fig. **S4**), using also the wingspan obtained from the μCT reconstruction of our reference specimen (Fig. **S5**). This procedure yields an effective vein density of 0.7 g cm^−3^, which is adopted for all subsequent analyses. Details are provided in the Supplementary Material (§S1.3).

With this density estimate, the pixel-wise characterisation of wing inertia (Eqn 1) is complete. For two specific cases within subsequent analysis (Fig. **9**B & Fig. **10**D), we use the resulting area ratio–inertia ratio relationship for *D. melanogaster* to derive damage area ratios from damage inertia ratios across Diptera. This assumed generalisation is motivated by (**i**) the conserved venation pattern across Diptera, and (**ii**) the fact that the normalisation of this ratio relationship means that it is exactly valid for any combination of chordwise, spanwise, or thin-plate depthwise stretching applied to the *D. melanogaster* wing—sufficient to describe the wing morphology of many other Dipterans. For comparison, we also construct also two lower-fidelity models, involving (**i**) the assumption that the surface density of the wing is uniform— a uniform plate model, as per Combes and Daniel (2003); and (**ii**) the assumption that the wing is a rectangular prism with uniform density—a uniform rectangular prism model (Salem et al., 2022). Details of these lower-fidelity models are given in the Supplementary Material (§S1.4).

### 2.2 Effective mass and inertia of thoracic deformation

During operation of the flight motor, the vibration of the thoracic structure necessarily entails some effective mass—or, in terms of wingbeat coordinates, some effective rotational inertia, *cf*. Agrawal et al. (2025). To estimate these effective properties, we must estimate (**i**) the pattern of deformation within the thorax, and (**ii**) the amplitude of this deformation—determining the transmission ratio between thoracic vibration and wingbeat. Representative deformation amplitudes can be derived from several sources, including muscle strain estimates (Chan and Dickinson, 1996; Mokso et al., 2015), but determining the thoracic deformation pattern is more challenging. To approximate this pattern, we develop an anatomically-realistic finite-element (FE) model of the thorax based on μCT data for *D. melanogaster* (Maisano, 2024).

Using 3D Slicer, MeshMixer, and Blender, we reconstruct a three-dimensional FE model of the thorax from this μCT data, accounting for the major thoracic sclerites (Fig. **3**B). Details of the reconstruction procedure and FE modelling are provided in the Supplementary Material (§S2). FE analysis is carried out in COMSOL Multiphysics, with the condition of sagittal symmetry: the thorax is modelled as a solid domain and the sclerites as shell elements coupled through a thin-to-solid interface (Fig. **3**C). To estimate representative thoracic vibration patterns, we perform a modal analysis: we anticipate that real *in vivo* deformation is some composition of the underlying thoracic modeshapes, and so FE estimates of the effective masses of these modeshapes provides insight into the effective mass of the *in vivo* pattern. A mesh convergence study confirms that our selected mesh is convergent (Supplementary Material, Fig. **S6**).

**Fig. 3.**
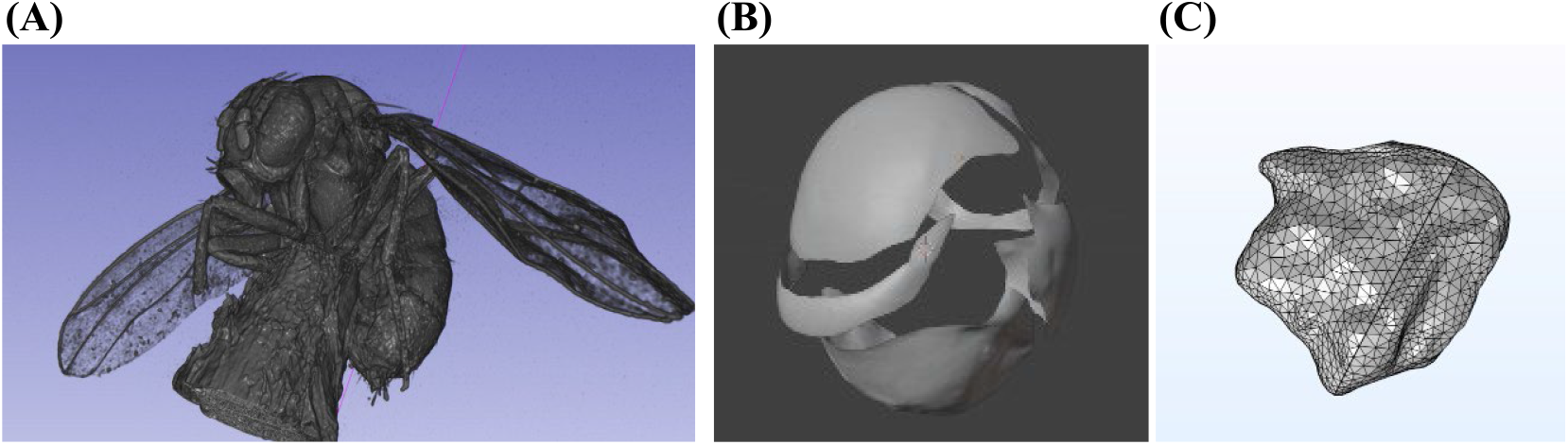
Workflow for thoracic reconstruction and finite-element (FE) modal analysis. (**A**) *D. melanogaster* reconstructed from μCT data (Maisano, 2024). (**B**) Refined geometry of the major sclerites, prepared in Blender prior to export into COMSOL. (**C**) Sagittally-symmetric FE mesh model used for modal analysis: the thorax is modelled as a solid domain and the sclerites as shell elements coupled through a thin-to-solid interface.

In the context of thoracic deformation, we define the parameter:

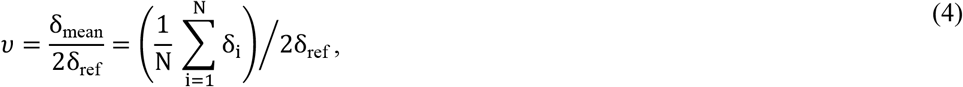

where *δ*_*i*_ represents the Euclidean displacement field of the structure, evaluated at the mesh Gaussian points (*i*): *δ*_mean_ is the mean displacement over the structure, and *δ*_ref_ the displacement at the top of the scutum, as a reference point. *v* thus, indicates how well scutal displacements (*i*.*e*., approximately, the DVM/DLM strains) represent the thoracic motion as a whole: were the entire thorax moving as a rigid body (a rigid-body mode), *v* = 1, but we expect *v* < 1. The factor of two (2*δ*_ref_) accounts for the fact that in the modeshapes of a thoracic structure with free boundary conditions, the scutal displacement will be half that of with the sternum fixed.

Using *v*, we compute the effective mass (M_t, eff_) and rotational inertia (I_t, eff_) of the thorax as:

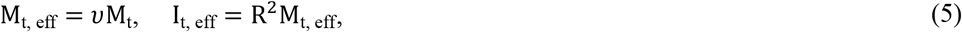

using the transmission ratio R, defining the ratio between thoracic deformation at a reference point and wingbeat stroke angle, see Eqn 13. To estimate *v*, we compute the first two modeshapes of the *D. melanogaster* thoracic FE model: these modeshapes are illustrated in Fig. **8**A. To assess the relative significance of thoracic effective mass vis-à-vis (intact) wing inertia, under classical PEA model, we define the parameter γ:

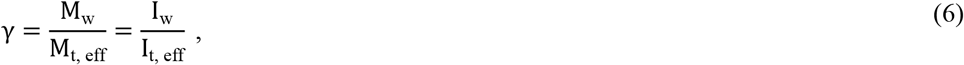

where large γ (≫ 1) indicates that the wing inertia is dominant. Meta-analysing data from a range of dipterans (Belyaev et al., 2012; Chan and Dickinson, 1996; Ennos, 1989; Lehmann and Dickinson, 1997; Marden, 1987; Marden, 2000; Martín-Vega et al., 2017; Peckham et al., 1990; Soares et al., 2015; Sotavalta, 1952), we predict γ across these species and the associated scaling of wingbeat frequency with wing damage, using the PEA modelling with constant motor stiffness (§2.4.1).

### 2.3 Viscoelasticity of activated flight muscle

Activated asynchronous flight muscle may be represented as a viscoelastic material (Maughan et al., 1998; Swank, 2012). At low strain amplitudes, stretch-activation manifests as a frequency-dependent complex modulus, E(Ω):

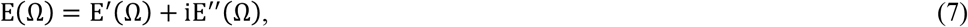

where E^′^(Ω) and E^′′^(Ω) denote the storage and loss moduli, respectively. The storage modulus reflects the elastic behaviour of the muscle, *i*.*e*., its effective spring-like stiffness. The loss modulus, and associated negative loss tangent r(Ω) = −E^′′^(Ω)/E^′^(Ω), reflect the energy the muscle outputs (or dissipates). Complex moduli data for ten indirect flight muscle fibres of *D. melanogaster* are reported in the dataset of Viswanathan et al. (2020), who, additionally, provide fitted moduli curves of the form (Swank, 2012):

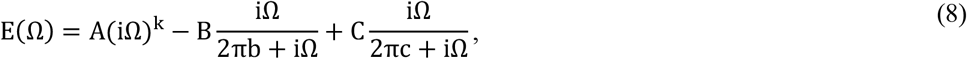

with constants A, B, C and k, b, c. In this dataset we observe that, over the frequency range that the muscle does work (Ω : r(Ω) > 0), the storage modulus shows an approximately linear decreasing trend. Fig. 4 illustrates this trend in normalised terms: at the upper frequency limit of work output, the storage modulus is less than 40% of its value at the lower limit.

**Fig. 4.**
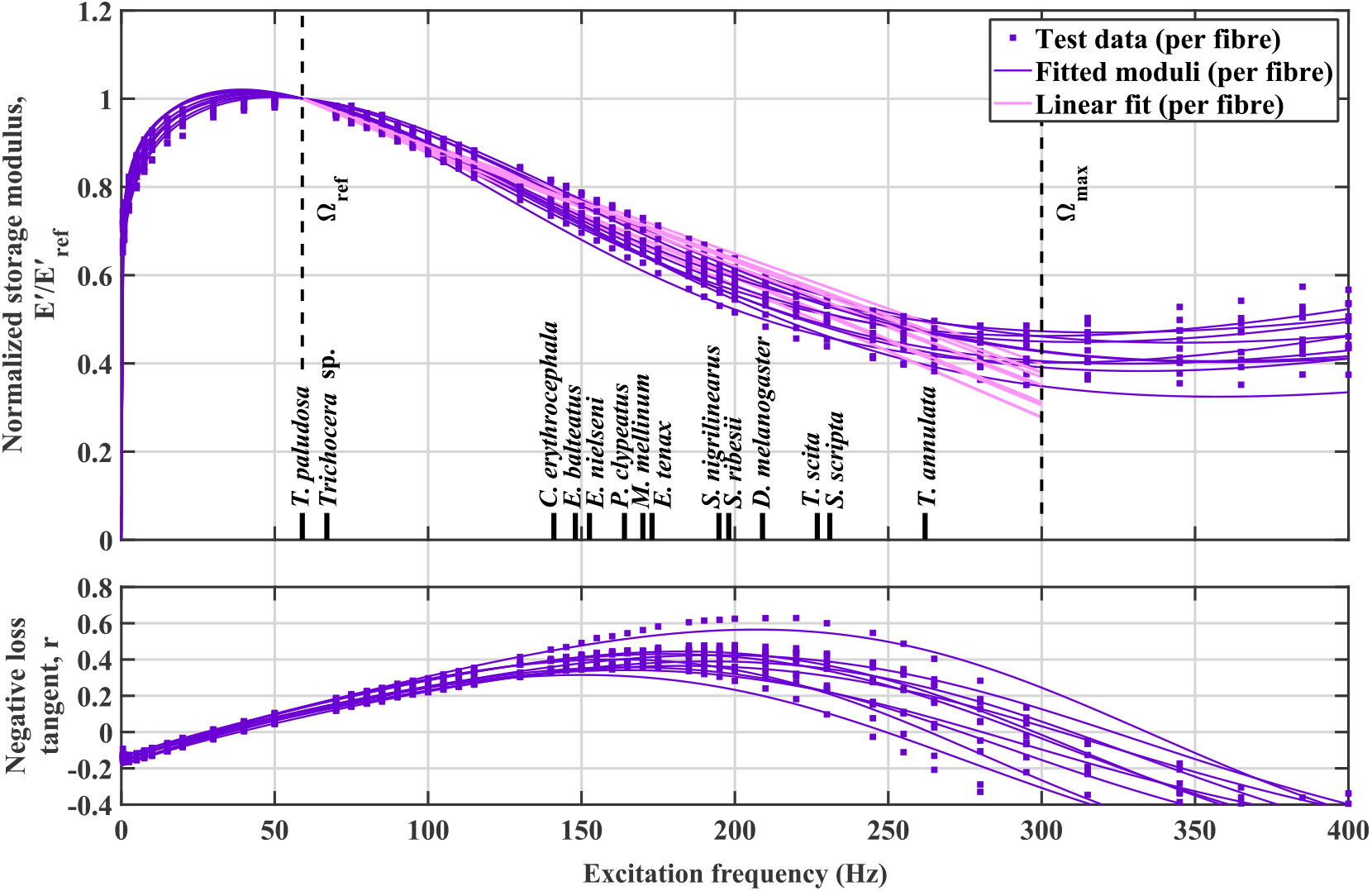
Viscoelastic characterisation of activated *D. melanogaster* flight muscle. (**A**) Trend in normalised storage modulus, E′(Ω)/E^′^(Ω_ref_), based on experimental data and fitted moduli curves from Viswanathan et al. (2020). Vertical markers indicate wingbeat frequencies of a range of dipterans: in addition to species in Fig. **1**, the hoverflies *Eupeodes nielseni, Tropidia scita, Sphaerophoria scripta, Syrphus nigrilinearus, Melanostoma mellinum*, and *Platycheirus clypeatus* (Belyaev et al., 2012; Ennos, 1989; Sotavalta, 1952). To illustrate a maximal effect, we take *Ω*_ref_ as the lowest wingbeat frequency among these species: that of the crane fly *T. paludosa*. (**B**) Corresponding negative loss tangent, r(Ω): where r > 0, the muscle does work.

To this data and trend, we compare the average wingbeat frequencies of a range of dipteran species (Fig. **4**)—specifically, those studied by Sotavalta (1952). We note that there is a broad open question of whether drosophilid muscle is representative of Diptera more widely: these meta-analysed wingbeat frequencies do lie within the work-output region, and so could hypothetically be generated by drosophilid muscle, but it may be that timescales of stretch-activation in these different species are adapted so that the wingbeat frequency lies closer to the frequency and which work output is maximised. Being unable to resolve this open question in this work, we aim instead to characterise the maximal storage modulus trend: an upper bound on the impact of storage modulus trends on wingbeat frequency scaling with damage.

We use the lowest wingbeat frequency reported by Sotavalta (1952), for the crane fly *Tipula paludosa*, as a reference frequency (Ω_ref_), and, over Ω_ref_ ≤ Ω ≤ Ω_max_, we perform a linear regression on the fitted fibre profiles. This leads to the piece-wise linear approximation:

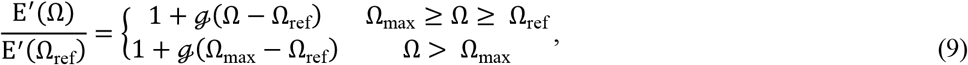

where ℊ is the gradient of the normalised trend, and above Ω_max_, the storage modulus is approximated as constant, following the observed trend in moduli data (Fig. **4**). We estimate ℊ = (−4.36 ± 0.29) × 10^−4^ across the ten muscle fibres.

This estimate corresponds to a maximal effect, *i*.*e*., most extreme loss of muscle stiffness, corresponding the case where *T. paludosa* has muscle properties identical to those of *D. melanogaster*. From this maximal estimate, we may approximate other submaximal effects associated with different cases. *E*.*g*., *D. melanogaster* itself is expected show only ≈30% of maximal effect; *C. erythrocephala*, were it to show muscle properties identical to of *D. melanogaster*, would show ≈70% of maximal effect; and so on.

### 2.4 Candidate models of the flight motor

#### 2.4.1 Parallel elastic actuation (PEA)

To explore the effect of the distribution of elasticity within the flight motor on wingbeat frequency scaling with damage, we evaluate several candidate models of the flight motor. First, we consider the classical linear single-degree-of-freedom (1DOF) PEA model, as in Fig. **5**A (Gau et al., 2022; Lynch et al., 2021; Pons and Beatus, 2022). In this model the wing and thorax are rigidly linked, and so we may define a transmission ratio, R, between wing stroke angle (θ, units: rad) and a reference thoracic displacement (u, units: m) as R = u_max_ /θ_max_. Using this transmission ratio we can define a translational proxy variable for the wing stroke angle, x = Rθ (units: m) and translational equations of motion:

**Fig. 5.**
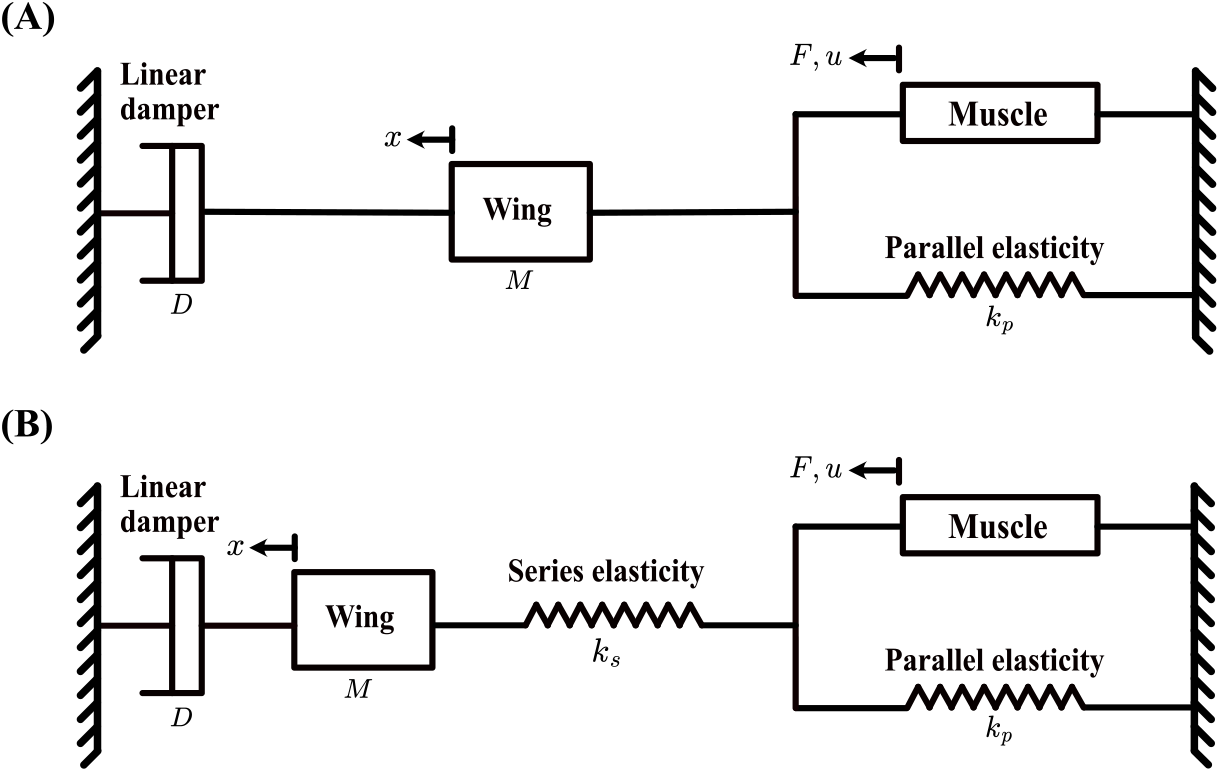
Candidate mechanical models of the dipteran flight motor. (**A**) Linear one-degree-of-freedom (1DOF) parallel elastic actuation (PEA) model, in which muscle actuation and thoracic elasticity (k_p_) act in parallel, and are rigidly coupled to the wing mass (M) and damping (D). (**B**) Linear 1DOF hybrid elastic actuation (HEA) model, incorporating both parallel thoracic elasticity (k_p_) and an additional series elastic element (k_s_) representing flexion at the wing root: this flexion modulates the coupling between the thorax and wing.

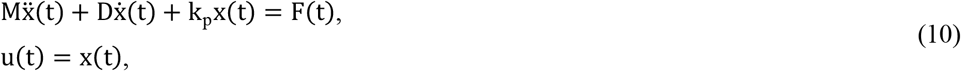

where M is the total effective translational mass of the motor (wing and thorax); D is the total damping coefficient of the motor—assuming linear damping; and k_p_ is the total stiffness of the motor (exoskeleton and muscle). In canonical form, Eqn 10 becomes:

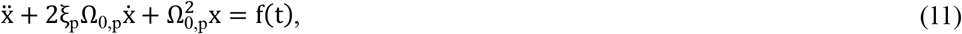

with parallel natural frequency Ω_0,p_ and linear damping ratio ξ_p_ defined as:

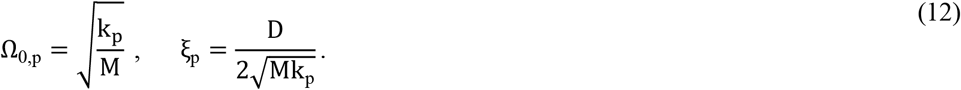

Via the transmission ratio R, we may compile wing and thorax inertial properties. Conserving wing kinetic energy across translational and rotational coordinates, 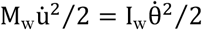, yields:

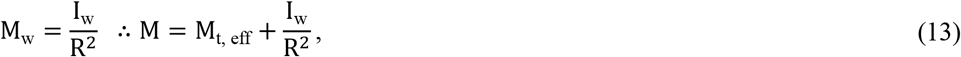

In advance of the results of §3.2, we note that thoracic effective inertia is not significant across the dipterans we study, and so for much of our further analysis, M = M_w_.

Within this motor model, we consider two possible assumptions regarding the properties of the primary flight muscles. *First*, we assume constant motor stiffness k_p_: this assumption would be valid if either the muscle complex modulus is constant or a linear thoracic elasticity dominates this complex modulus. By self-oscillation to effective energy resonance (Pons, 2023), the frequency to which the wingbeat will converge is Ω_0,p_, and thus the scaling of the wingbeat frequency with wing inertia ratio is:

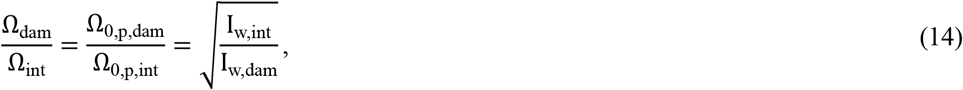

equivalent to the classical result of Salem et al. (2022). However, as per Fig. **4**, the storage modulus of *D. melanogaster* flight muscle is not constant; and evidence suggests that this modulus is the dominant stiffness in the *D. melanogaster* flight muscle (Pons et al., 2023). As a *second* assumption, we take the piecewise storage modulus trend of Eqn 9 to represent k_p_, *i*.*e*., that this maximal trend represents the complete trend in motor effective stiffness. Thus:

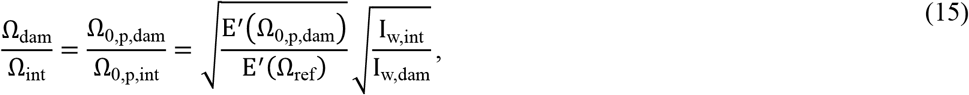

which is an implicit relation: E^′^ depends itself on Ω_0,p,dam_. We solve this implicit relation for the case of the linear storage modulus trend, Eqn 9, to yield:

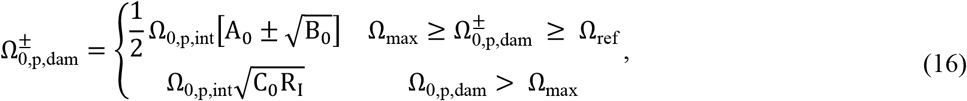

where the auxiliary parameters R_I_, A_0_, B_0_, C_0_ are defined as:

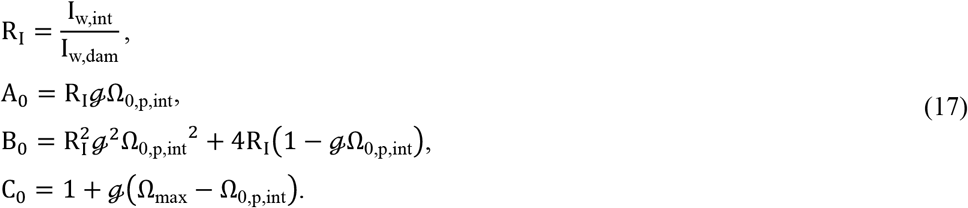

This scenario represents a depressed scaling w.r.t. to the classical result (Eqn 14).

#### 2.4.2 Hybrid elastic actuation (HEA) with wing root flexion

Following Wang et al. (2026), we consider a second candidate motor, with the addition of a flexural (series) element at the wing root—a case of hybrid-elastic actuation (HEA), leading to phase lags between the wingbeat and muscle. We note that there is further precedent for such a model in Diptera: Melis et al. (2024) observed wing-root flexure in *Drosophila hydei* corresponding to an phase difference of approximately 0.28 rad. A linear 1DOF HEA model with wing root flexion (Fig. **5**B) may be expressed:

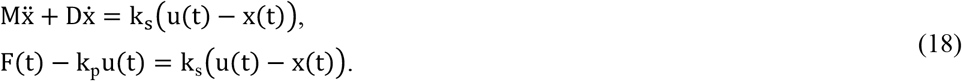

In addition to parameters from the PEA model, this HEA model contains a series elasticity k_s_, defining a natural frequency (Ω_0,s_), damping ratio (ξ_s_) and elasticity ratio (α):

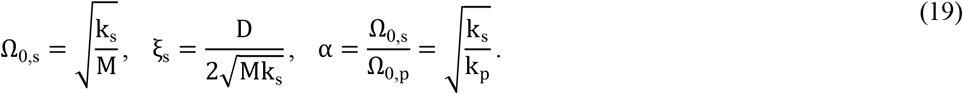

Note that ξ_s_ and ξ_p_ are not unique: they are merely rescalings of D in series and parallel reference frames. The parameter α quantifies the relative strength of the series elasticity with respect to the parallel elasticity. Finite values of α represent any system that is neither pure PEA (Ω_0,s_ → ∞), nor pure SEA (Ω_0,p_ = 0).

The self-oscillation frequency of this HEA model has not previously been presented. A derivation of this self-oscillation frequency, along with numerical validation is provided, in the Supplementary Material (§§S4.1, S4.2). We compute it as:

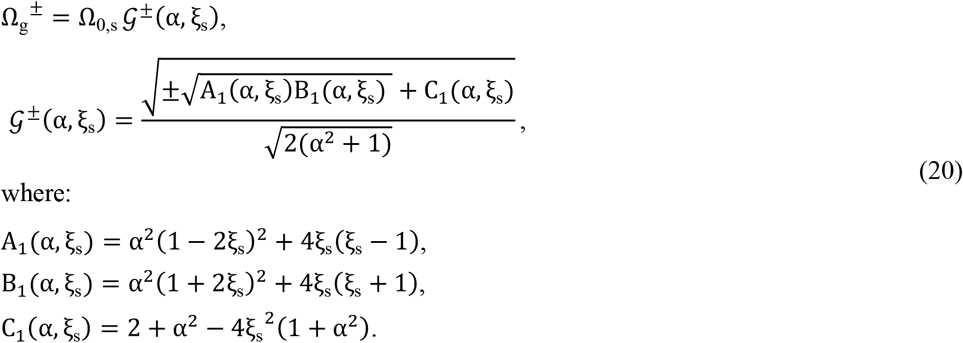

Here there are two solutions: 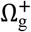 and 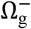 corresponding to SEA-like (wing hinge-dominated) and PEA-like (thorax-dominated) resonance, respectively: we consider only the PEA-like solution 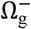. Eqn 20 has a notable structure: Ω_g_^±^ scales linearly with Ω_0,s_, with all dependency on α and ξ_s_ encoded within *G*^±^(α, ξ_s_).

#### 2.4.3 Scaling of linear damping ratio with wing damage

A broad open question is how the effective linear damping ratio of the motor (ξ_s_ or ξ_p_) scales with the level of damage to the wing. In the case of the PEA models, Eqn 14-17, the self-oscillation frequency is independent of the damping ratio, and so it is not necessary to estimate this scaling. However, the presence of wing root flexion in the HEA model breaks this independence, and ξ_s_ influences damage-frequency scaling via *G*^±^(α, ξ_s_). We derive an approximate model-based scaling relation for ξ_s_. Details of this derivation are given in the Supplementary Material (§S4.3). We approximate the wing as a rectangular plate with quasi-steady aerodynamic drag, with wing inertia scaling computed from the rectangular plate model (Salem et al., 2022). This yields the scaling relation:

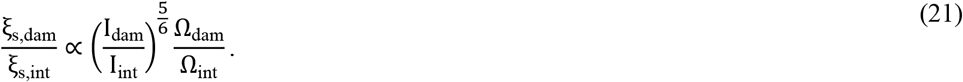

Incorporating Sotavalta’s (1952) empirical power-law scaling of frequency with damaged-wing inertia, we then derive the explicit relationship:

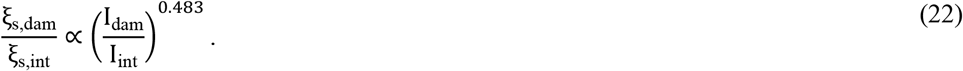

#### 2.4.4 Scaling of HEA motor self-oscillation frequency with wing damage

To compute the overall scaling of HEA motor self-oscillation frequency with wing damage, we may combine the motor dynamics, damping ratio, and muscular property scaling relations derived so far. First, under the assumption that the motor stiffness is constant (*i*.*e*., muscular storage modulus is constant or negligible), we may express:

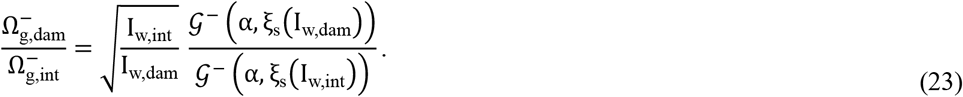

Second, under the assumption that the motor stiffness shows the maximal linear scaling associated with the muscle storage modulus (*i*.*e*., muscle stiffness dominates exoskeletal stiffness), we then have the additional effect that the stiffness ratio α is no longer constant. We must therefore define an α ratio term:

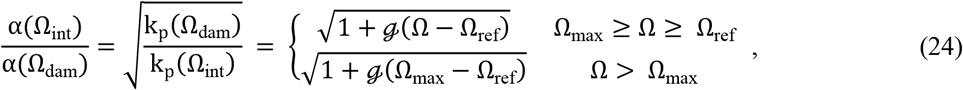

which decreases monotonically, from 1, until Ω_max_. Beyond Ω_max_, this ratio remains unchanged at its minimum value (*cf*. Fig. **4**). We may then express the frequency-damage scaling as:

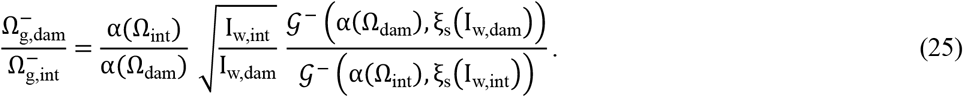

This scaling relationship is implicit, due to the dependency of α on frequency. To compute 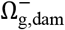 explicitly, we employ a numerical continuation method in which the inertia-ratio domain is discretised and the frequency-dependent stiffness is updated progressively based on the computed frequency at the previous step. Details of this numerical method are given in the Supplementary Material (§S4.4). Using it, we obtain frequency scaling profiles for a representative Dipteran motor with ξ_p,int_ = 0.29—illustrative, *e*.*g*., of *Tipula* sp. (ξ_p_ ≈ 0.11, Ellington, (1984)) and *D. hydei* (ξ_p_ ≈ 0.31, Dickinson and Lighton, (1995)), and for 1.4 ≤

α(Ω_int_) ≤ 3, indicative of wing flexion in drosophilids (Wang et al., 2026). We note that, to guarantee that *G*^−^(α, ξ_s_) remains real-valued (*i*.*e*., self-oscillation occurs), a boundary over ξ_p_ and α exists—we analyse this boundary in the Supplementary Material (§S4.4).

## 3 Results

### 3.1 Drosophilid wing inertia is strongly non-uniform over wingspan

Previous approaches to model dipteran wing rotational inertia—whether, for damage scaling (Salem et al., 2022), or for general aeromechanical characterisation (Combes and Daniel, 2003; Kolomenskiy et al., 2019; Pons et al., 2023; Zhao and Deng, 2009)—have assumed the wing mass to be distributed uniformly. However, our pixel-based reconstruction of the *D. melanogaster* wing predicts strongly non-uniform distributions of mass and rotational inertia (Fig. **6**). We observe two notable features of these distributions:

**Fig. 6.**
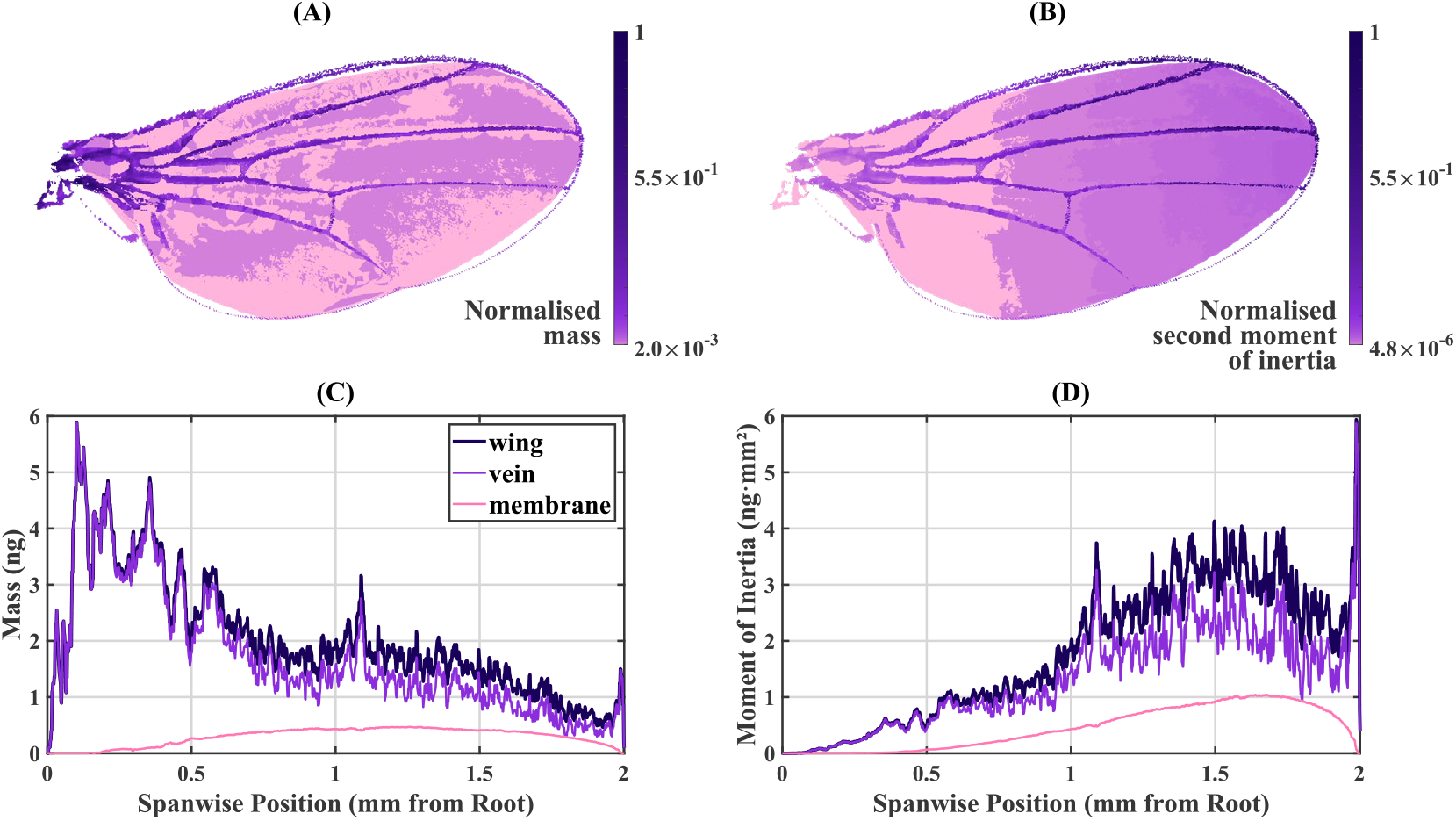
Spatial distributions of *D. melanogaster* wing inertia. **(A)** Map of normalised mass, where the mass at each pixel is scaled by the maximum mass over the wing. **(B)** Map of the normalised second moment of inertia about the wing root, where the inertia at each pixel is scaled by the maximum inertia. **(C)** Spanwise distribution of mass, decomposed by vein and membrane. **(D)** Spanwise distribution of the second moment of inertia, similarly decomposed.

**(i)** The mass and rotational inertia of the wing veins dominates those of the wing membrane: the veins account for 86% of the total mass and 75% of the total rotational inertia. An assumption that the wing mass is concentrated in the veins is thus more accurate than an assumption that it is distributed uniformly over the wing. Interestingly, this indicates that inertial properties of Dipteran wings may effectively be encoded in their vein architecture— and that changes in venation, whether through developmental variation or evolutionary modification, drive inertial changes and thereby motor mechanical ones.

**(ii)** The wing mass is concentrated within basal vein stems near the wing root—consistent with observations of similar concentration in bees (Kolomenskiy et al., 2019). However, these stems do not appreciably contribute to rotational inertia, which instead is dominated by the distal tips of the longitudinal veins L1-L4. In particular, the segment of the costa (L1) that lies between the tips of L2-L4 accounts for a large peak in spanwise inertia distribution.

From modelling perspective, conventional uniform plate and rectangular prism models lead to significant errors in the distribution of mass and rotational inertia (Fig. **7**A-B), ultimately leading to errors in scaling predictions based on area—Fig. **7**C illustrates the example case of unilateral wing clipping in a PEA motor with constant motor stiffness (Eqn (14).

**Fig. 7.**
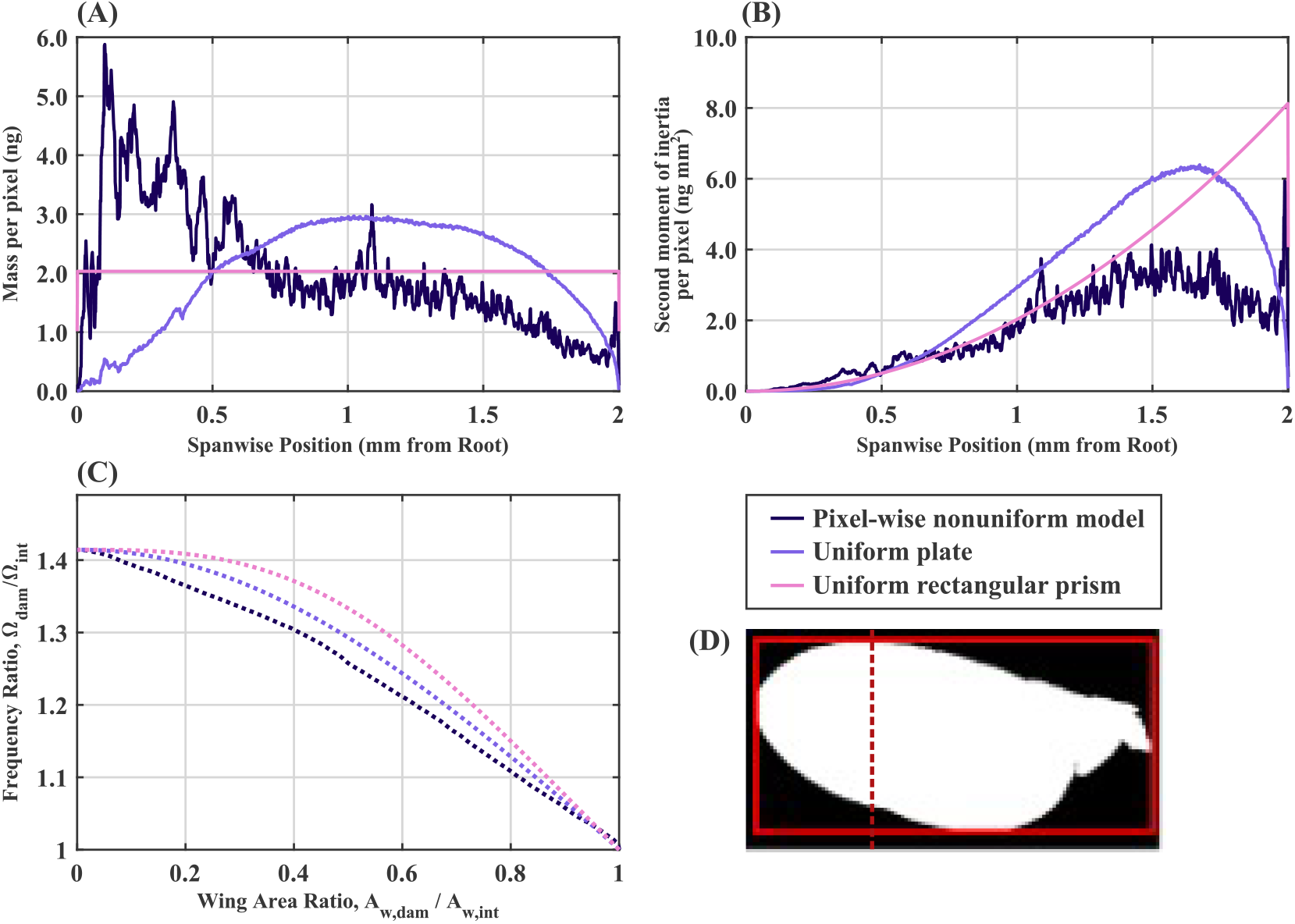
Model fidelity for a wing of D. melanogaster under different inertial assumptions. **(A)** Spanwise mass distribution, showing results from the pixel-based nonuniform model, uniform plate model, and uniform rectangular prism model. **(B)** Spanwise distribution of second moment of inertia about the wing root for the same set of models. **(C)** Predicted scaling of self-oscillation frequency with the ratio of damaged wing area, for the set of models, under unilateral chordwise wing clipping as illustrated in (**D**). All compared models have the same total wing mass.

### 3.2 Thoracic effective inertia is negligible across Diptera

Fig. **8**A illustrates the first two modes of the *D. melanogaster* thorax, computed by finite-element modal analysis. For both modes, the deformation metric *v* indicates that the effective thoracic mass is approximately 39% total thoracic mass. We find, as a result, that estimates of effective-mass ratio, γ, for many of the dipteran species considered in this study are very large: ranging between 10^2^ and 10^6^. This indicates that wing inertia dominates thoracic inertia across Diptera. To confirm that thoracic inertia still does not influence damage scaling, we incorporate these γ estimates into the PEA motor predictions: this produces frequency scaling curves that are visually indistinguishable from one another (Fig. **8**B). This indicates that, while dipteran species differ significantly in body size, thoracic effective mass has a negligible contribution to motor dynamics and associated scaling with wing damage. Interestingly, comparing these scaling predictions with the frequencies of non-flight vibration (defence buzzing) behaviour in hoverflies from genera Eristalis and Syrphus (Vallejo-Marín and Vallejo, 2021) indicates that another non-inertial mechanism must govern motor dynamics under these behaviours. With the wings retracted, wing inertia should be negligible under defence buzzing, but defence buzzing frequencies do not match the strong scaling of frequency predicted from inertial loss (Fig. **8**B).

**Fig. 8.**
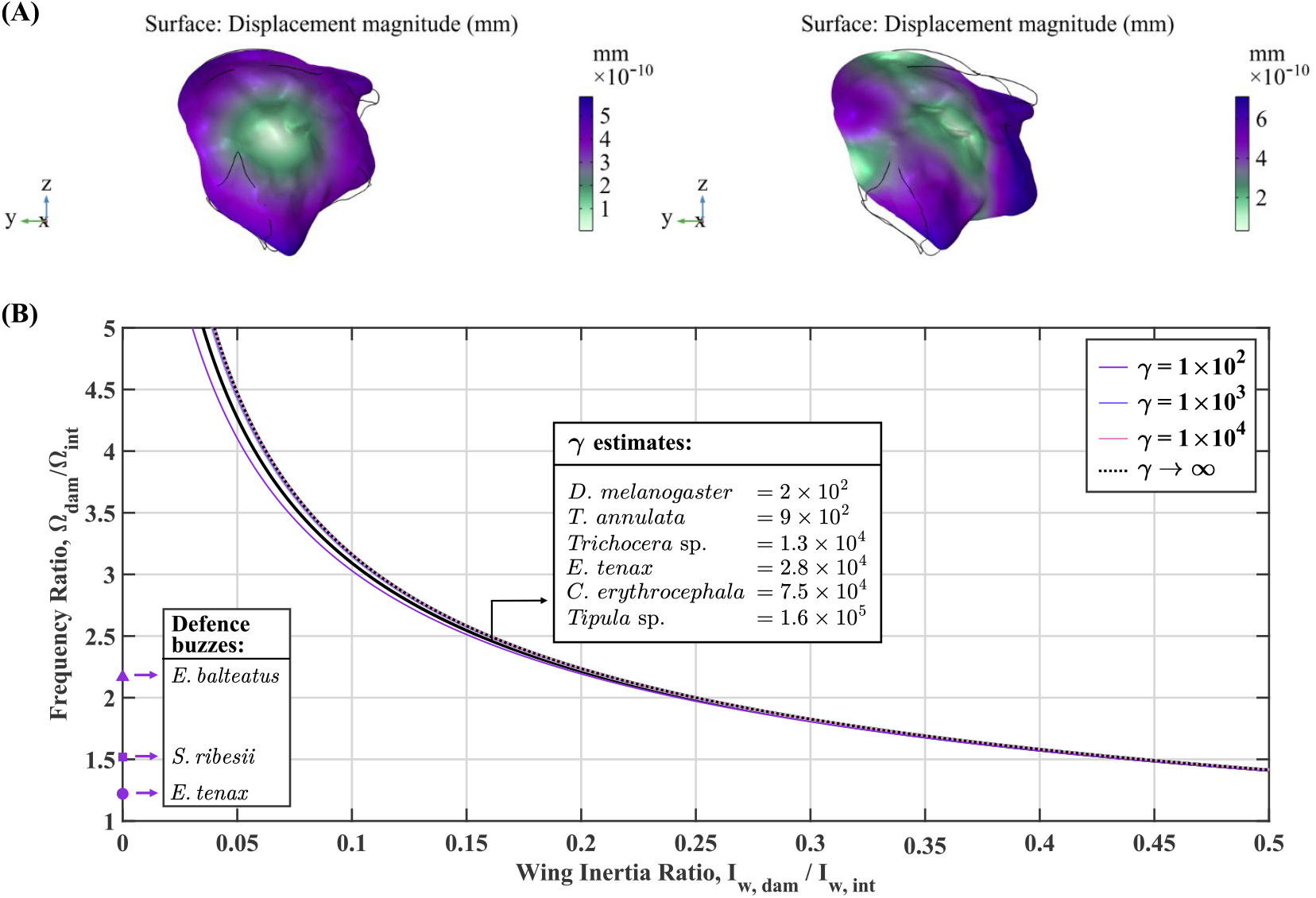
Negligible effects of thoracic effective mass across Diptera. **(A)** Displacement fields for the first two modes of the *D. melanogaster* thorax, computed by FE analysis. They are used to estimate the contribution of thoracic deformation to total flight motor inertia during motor operation. **(B)** PEA model predictions of self-oscillation frequency under wing damage, for constant muscle stiffness and combined wing-thorax inertia with varying *γ* = M_w,int_/M_t, eff_. Indicated are estimated values of *γ* across Diptera, based on meta-analysed kinematic and morphological data (Belyaev et al., 2012; Ennos, 1989; Marden, 1987; Marden, 2000; Martín-Vega et al., 2017; Soares et al., 2015; Sotavalta, 1952). Frequency-ratio measurements from hoverflies over defence buzzes (Vallejo-Marín and Vallejo, 2021) are included for comparison.

### 3.3 Parallel elasticities are unable to completely explain depressed frequency-damage scaling

Fig. **9**A illustrates the damage-frequency scaling predictions of PEA models—with and without the complex-modulus trend of flight muscle—overlaid against experimental data for bilateral wing clipping (Sotavalta, 1952) and wing folding during defensive buzzing (Vallejo-Marín and Vallejo, 2021). While these PEA models reflect the general upwards trend of this data, the constant-stiffness model markedly overpredicts the increase in wingbeat frequency. The model with the maximal effect of muscle complex modulus accounts partly, but not fully, for the depressed scaling observed in the experimental data, and we note that this effect is maximal: species-specific predictions would lie between these two models. And we observe again that errors in defence buzz predictions are particularly extreme. A similar effect is seen (Fig. **9**B) when comparing PEA model predictions to meta-analysed unilateral wing clipping data (Deora et al., 2021; Hao et al., 2025; Meng et al., 2023; Muijres et al., 2017; Salem et al., 2022; Sotavalta, 1953), associated with smaller changes in inertia and thus wingbeat frequency. Some of this data—for drosophilids—matches PEA model predictions on average, but much lies depressed of the PEA scaling curve. From this pair of results, we conclude that parallel elasticities do not themselves fully explain damage-frequency scaling.

**Fig. 9.**
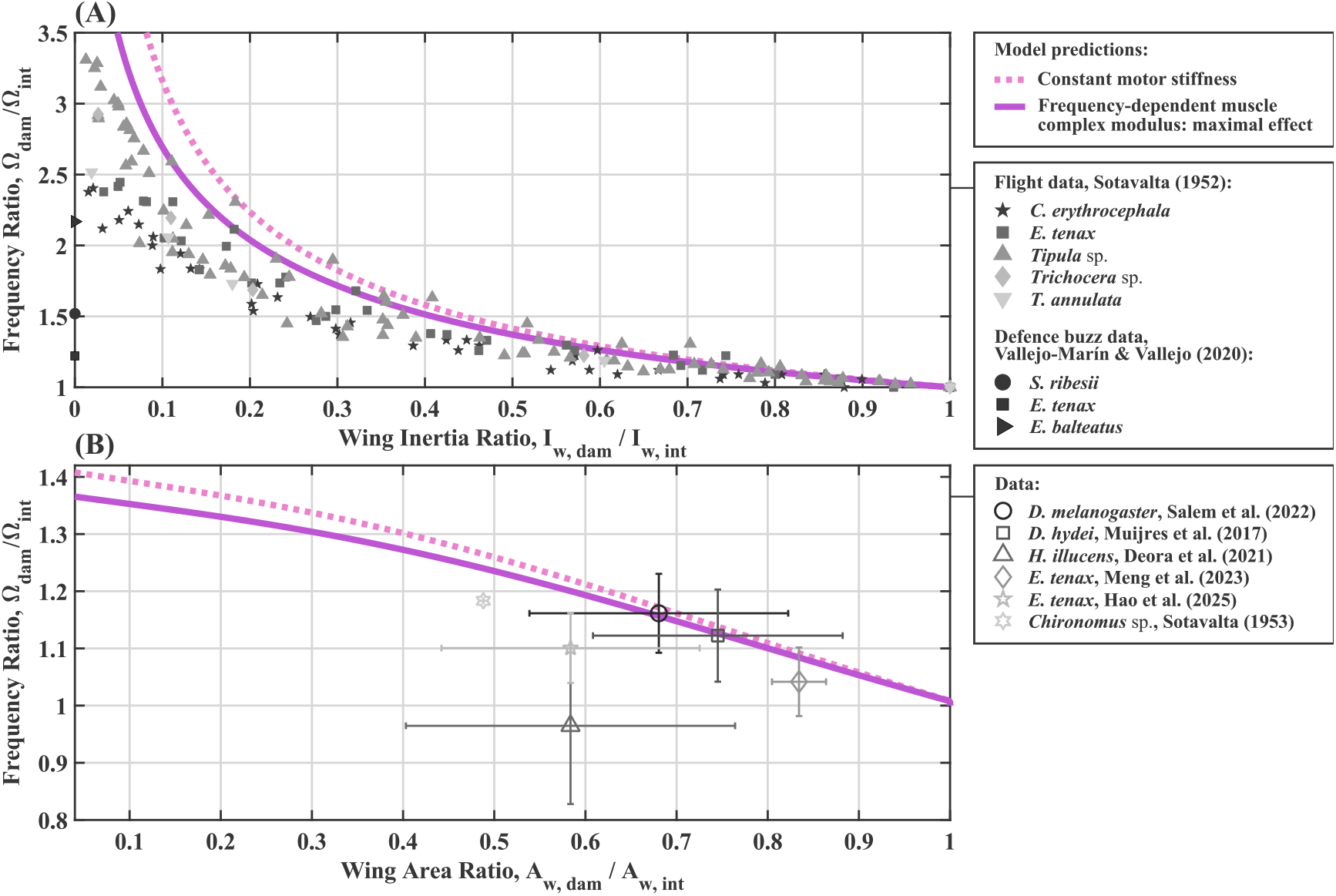
Comparison of PEA model predictions with experimental wing-clipping or wing-folding data. **(A)** Dipteran wingbeat frequency scaling under bilateral clipping or folding: wingbeat frequency ratio versus total (bilateral) wing inertia ratio for PEA models, overlaid on clipping data from Sotavalta (1952) and defensive buzzing data from Vallejo-Marín and Vallejo (2021). **(B)** Scaling under unilateral wing clipping: wingbeat frequency ratio versus the ratio of unilateral remaining wing area. PEA model predictions are obtained by converting wing area ratio to wing inertia ratio using a polynomial fit derived from the pixel-wise nonuniform model of the *D. melanogaster* wing. Meta-analysed experimental data for unilateral wing-clipping (Deora et al., 2021; Hao et al., 2025; Meng et al., 2023; Muijres et al., 2017; Salem et al., 2022; Sotavalta, 1953) are shown as mean±SD for comparison. Details of the meta-analysed data processing are provided in the Supplementary Material (§S3).

### 3.4 Wing root flexion depresses damage-frequency scaling

The presence of a series-elastic element at the wing root—within an HEA motor model— changes this picture substantially. Lower values of α, corresponding to a more compliant wing root, depress the scaling of wingbeat frequency with wing inertia under damage. This depressed scaling occurs even under constant motor stiffness (Fig. **10**A), solely due to the scaling of damping ratio with damage: in this scenario α remains constant under damage. However, the frequency-dependency of muscle complex modulus compounds this depression (Fig. **10**B): the decreasing complex modulus alters α which leads to a further decrease via *G*^±^(α, ξ_s_). Motor models with realistic values of α agree substantially better with experimental data for bilateral wing clipping (Fig. **10**C) as well as unilateral wing clipping (Fig. **10**D). We illustrate this improved agreement via a species-specific fit of α to data for two species reported by Sotavalta (1952): *C. erythrocephala* and an unidentified *Tipula* sp., on the basis of the error between predicted and observed frequency scaling (Fig. **10**E).

**Fig. 10.**
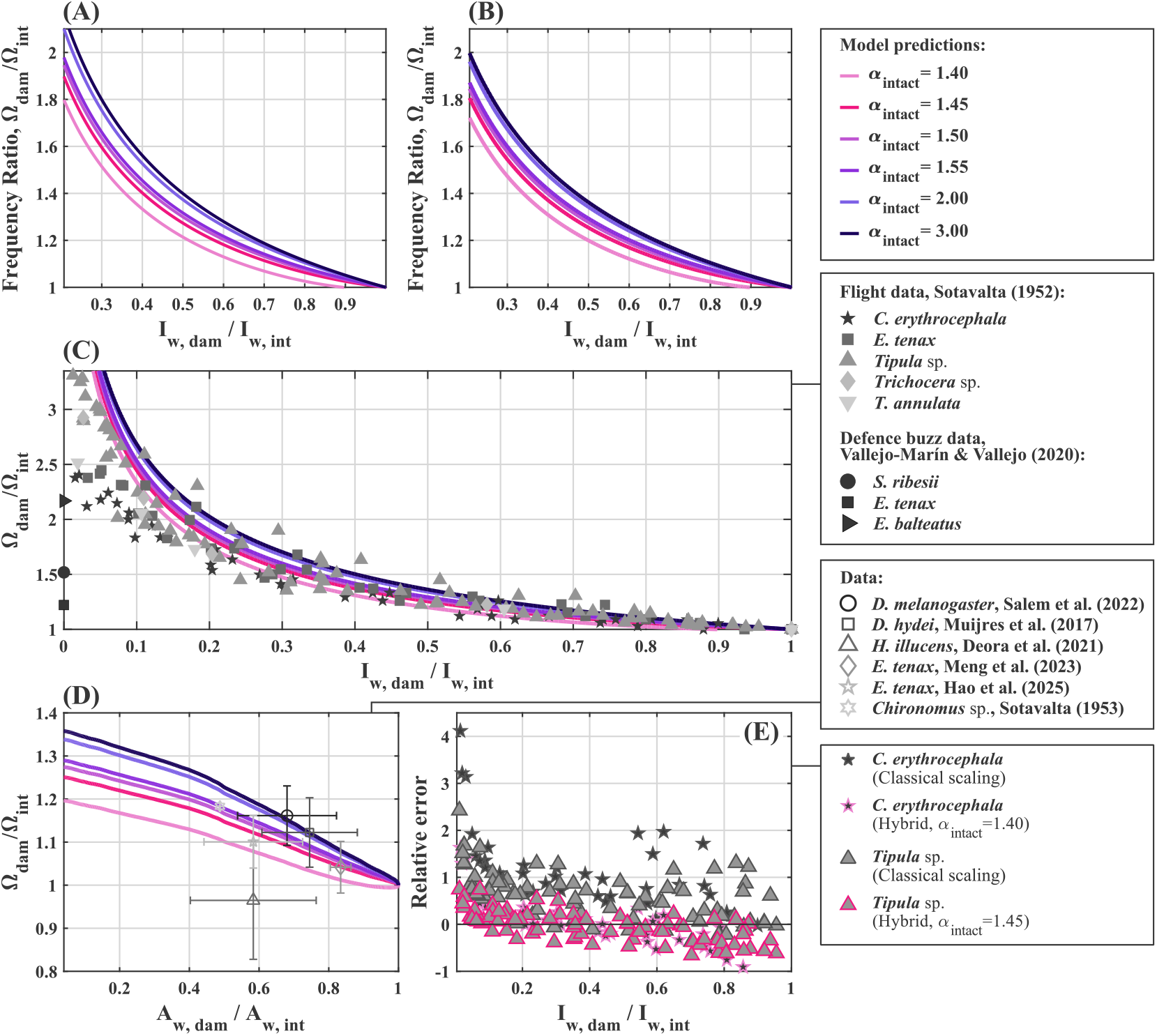
Responses of the hybrid (HEA) model to wing damage. **(A)** Scaling of wingbeat frequency with damage for an HEA model (*ξ*_*p*_ = 0.29) under constant motor stiffness, and **(B)** under the maximal decreasing trend muscle complex modulus. The case α_intact_ = α(Ω_int_) = 3 recovers PEA-like behaviour in either case. **(C)** Comparison of HEA model predictions under maximal-trend complex modulus with the bilateral clipping data of Sotavalta (1952) and the defensive buzz data of Vallejo-Marín and Vallejo (2021). **(D)** Comparison of HEA model predictions under maximal-trend complex modulus with meta-analysed data for unilateral wing clipping (Deora et al., 2021; Hao et al., 2025; Meng et al., 2023; Muijres et al., 2017; Salem et al., 2022; Sotavalta, 1953). To make comparison in terms of remaining wing area ratio, wing inertia ratio is converted via a polynomial fit derived from the pixel-wise nonuniform model of the *D. melanogaster* wing. **(E)** Quantitative illustration of the improvement in prediction: relative prediction errors, for two species reported by Sotavalta (1952): *C. erythrocephala* and an unidentified *Tipula* sp., comparing the classical scaling model with a species-specific best fit of α_intact_. The best-fit α_intact_ is similar for both species (≈ 1.4) and matches estimations of this structural property ratio for drosophilids (Wang et al., 2026).

Previous studies have attributed flight motor responses under wing damage to active sensory feedback (Bartussek et al., 2013; Deora et al., 2021; Duistermars et al., 2007; Muijres et al., 2017); in contrast, we suggest that much of this response arises passively from the motor’s mechanics. Taken together, our results indicate that depressed wingbeat-frequency scaling observed across Diptera likely arises not from any single passive feature in isolation, but from the interaction between wing root compliance, the viscoelasticity of the primary flight muscle, and the aerodynamic changes associated with progressive wing loss—and, not from thoracic effective inertia. The non-uniformity of wing inertia contributes to depressed scaling in terms of wing area, but not in terms of wing inertia itself. However, even our improved model, accounting for these interactions, cannot capture experimental observations at extreme levels of wing damage (<10% remaining inertia), and does not predict defensive-buzz frequencies. Both these features suggest that a further mechanism is at work in these extremal cases.

## 4 Discussion and Conclusion

### 4.1 Utility of spatially resolving wing inertia distribution

In this work we presented a general method to resolve the spatial distribution of mass and inertia over an insect wing—one which may have broader utility. This method draws largely upon visible-light imagery that is easily collected—for overall pixelwise classification (vein/membrane) and for membrane thickness estimation via WIP—but does require μCT data for vein thickness estimation, which is more involved to collect. It may be possible to replace this μCT dependency by estimating vein depth from vein width in visible-light imagery, and thereby develop an approach requiring only conventional optical microscopy imagery, but such an approach requires validation.

Regardless, this method may have wider utility in studies of the insect flight motor. Our results in *D. melanogaster*—that, vein inertia dominates membrane inertia—suggest that adapting venation patterns could act as a mechanism to adapt wing inertia independent of any significant aerodynamic effects. The morphology and phylogeny of venation is already well-studied (Pass, 2018; Perrard et al., 2016; Sharkey and Roy, 2002; Stark et al., 1999; Sugiyama et al., 2024): resolving the associated inertia distributions might offer further insights into the pressures driving the diversification and or dimorphism of vein patterns. There is also the prospect of applying this method to natural patterns of wing wear to assess their impact on the flight motor. The nonuniformity of the distribution of inertia over the wing indicates that the inertial effects of damage will differ markedly by the location of damage: not only, proximal *vs*. distal (as predicted by classical uniform-inertia models), but also leading-*vs*. trailing edge and whether damage affects the membrane or vein (a distinction invisible to uniform-inertia models). We conjecture that, since natural wear is liable to affect the membrane over the veins (Mountcastle and Combes, 2014; Rajabi et al., 2020; Rajabi et al., 2017), this wear is likely to have a greater aerodynamic impacts than inertial one—a prediction that would inform any assessment of the adaptive value of damage-frequency scaling, as we now discuss.

### 4.2 Does depressed damage-frequency scaling have any adaptive value?

A wide range of experimental data indicates that the scaling of wingbeat frequency with wing damage is depressed relative to classical parallel-elastic motor predictions (Fig. **1**). Accounting for wing root flexion, the viscoelasticity of the primary flight muscles, and the spatial nonuniformity of wing inertia recovers this depressed scaling in a motor model. However, the adaptive value or maladaptive impact (if any) of such scaling remains an open question. We conjecture that the evolution of insect wingbeat responses to wing damage is driven by two competing factors: (**i**) ensuring sufficient lift force to maintain flight with damaged wings— motivating increases to wingbeat frequency; and (**ii**) ensuring that the primary flight muscles can generate sufficient power and/or efficiency to drive the wingbeat—motivating a wingbeat frequency that remains close to a muscle optimum frequency (Fig. **4**).

These competing factors might, in general, drive evolution of a depressed scaling of wingbeat frequency with wing damage, though assessing this quantitively would require more data on the force generation properties of flight muscle across Diptera more widely—currently, such data is restricted to drosophilids. It would also require analyses of natural wing wear patterns rather than artificial wing clipping—our pixelwise method provides a basis for analysing such patterns (§4.1). However, it is also possible that the morphological features that drive this depressed scaling have unrelated function: there is evidence that wing root flexion may improve the efficiency of wingbeat frequency modulation during manoeuvring (Wang et al., 2026). In any case, depressed damage-frequency scaling is likely only evolutionarily significant at lower levels of wing damage, *i*.*e*., higher inertia ratio: at ratios below, *e*.*g*., 10%, the wings are effectively ablated and flight is likely impossible under any circumstances.

### 4.3 Prevalence of motor features across Diptera

Our explanation of depressed wingbeat scaling with damage hinges on the presence of a compliant wing hinge: via the hinge stiffness (α), aerodynamic damping and muscular viscoelasticity generate depressed scaling. The fact that depressed scaling is observed across a wide range of dipterans implies that that this wing hinge compliance might be similarly widespread—a picture consistent with limited existing observations in drosophilids (Melis et al., 2024) as well as other insect orders (Casey et al., 2025). However, we note also that our analysis focused on the presence of concentrated (lumped) flexion at the wing root, following Melis et al. (2024), rather than the alternative of distributed flexion along the length of the wing—such distributed flexion has been observed in other dipterans, such as *E. tenax* (Walker et al., 2009). Based on our analysis results for root flexion, it appears likely that distributed flexion would also lead to the depressed scaling we seek to explain, and so more broadly we would predict that either wing root flexion and/or wing flexion should be prevalent in Diptera. We conjecture that species with more slender, low-aspect-ratio, or weakly veined wings might rely more on distributed flexion than localised flexion.

We note also that these results are consistent with—though, do not directly prove—indications that the storage modulus of the primary flight muscles is the dominant stiffness within the dipteran flight motor, accounting for essential elastic energy storage, with exoskeletal elasticity contributing little (Pons et al., 2023; Alexander, 1982; Alexander and Bennet-Clark, 1977). In combination with wing root flexion, strong trends in decreasing motor stiffness with frequency play a significant role in explaining depressed wingbeat frequency scaling with damage. The limitations on direct proof in this regard are that it is possible that wing root and/or wing flexion that is more extreme than previously observed could explain the contribution that we attribute to muscular viscoelasticity; and that there remain unresolved inconsistencies regarding the stiffness of asynchronous muscles within the drosophilid motor (Pons, 2023). Nonetheless, it remains the case that depressed wingbeat frequency scaling with damage is consistent with the dominance of muscular stiffness; and, given the prevalence of depressed scaling in Diptera, this dominance is plausibly general to the order—suggesting the future comparative work might target the muscle complex modulus rather than thoracic material properties, *cf*. Casey et al. (2022); Deora et al. (2017) and Jankauski, (2020).

## Acknowledgements

We would like to thank Mario Vallejo-Marín for pointing us toward defence buzzing in hoverflies, as well as Chris Jones and Stephen De Lisle for providing original images from their work.

## Data availability

Code and data are attached as supplementary material.

