## Supplementary Material for "Dipteran wing damage discriminates between flight motor inertial-elastic distributions"

#### **S1 Pixel-wise reconstruction of nonuniform wing inertia**

##### **S1.1 Pixel-wise framework**

The optical image of wing selected for processing must be of high quality, as it directly influences the accuracy of subsequent feature extraction and segmentation. In this study, a high-resolution isotropic RGB image of a *D. melanogaster* wing is used as the input dataset, obtained from Flickr (Jones, 2013). The RGB wing image is pre-processed to suppress fine-scale membrane texture and enhance intensity differences between veins, membrane, and background prior to quantitative analysis, as shown in Fig. S1A. The blurred RGB image is then converted to a grayscale intensity image, isolating luminance information from colour, which is essential for identifying structural features such as veins and wing boundaries that are primarily defined by brightness variations rather than chromatic content. To enhance local contrast, we apply Contrast Limited Adaptive Histogram Equalization (CLAHE) to the grayscale image. This operation improves visibility of fine structures, particularly dark vein patterns and weak boundaries, while limiting noise amplification in homogeneous regions, as illustrated in Fig. S1B. Structural features are further emphasized through edge detection applied to the contrast-enhanced grayscale image. The resulting edge map highlights wing contours and internal vein boundaries, providing complementary information that is independent of color-based segmentation.

In parallel, color information from the original RGB image is incorporated through unsupervised clustering: pixel-wise k-means clustering is applied in RGB colour space to partition the image into a small number of representative regions corresponding to the membrane, veins, and background. Cluster identities are inferred based on relative intensity characteristics, with darker clusters associated with vein structures and brighter clusters associated with the surrounding background. To obtain a robust wing domain, the color-based segmentation is combined with the edge-derived structural information (Fig. S1C). Morphological refinement is subsequently applied to remove small artifacts, fill enclosed gaps, and retain only the largest connected component, which is assumed to correspond to the wing. Based on this refined wing mask, a labeled image is constructed in which pixels are assigned to one of three classes: vein, wing membrane, or outer background (Fig. S1D). The resulting labeled image provides a clear pixel-level representation of wing anatomy suitable for downstream geometric reconstruction and inertia estimation. This integrated processing framework combines intensity-based structural analysis with color-based region clustering to achieve reliable segmentation of biologically relevant wing features.

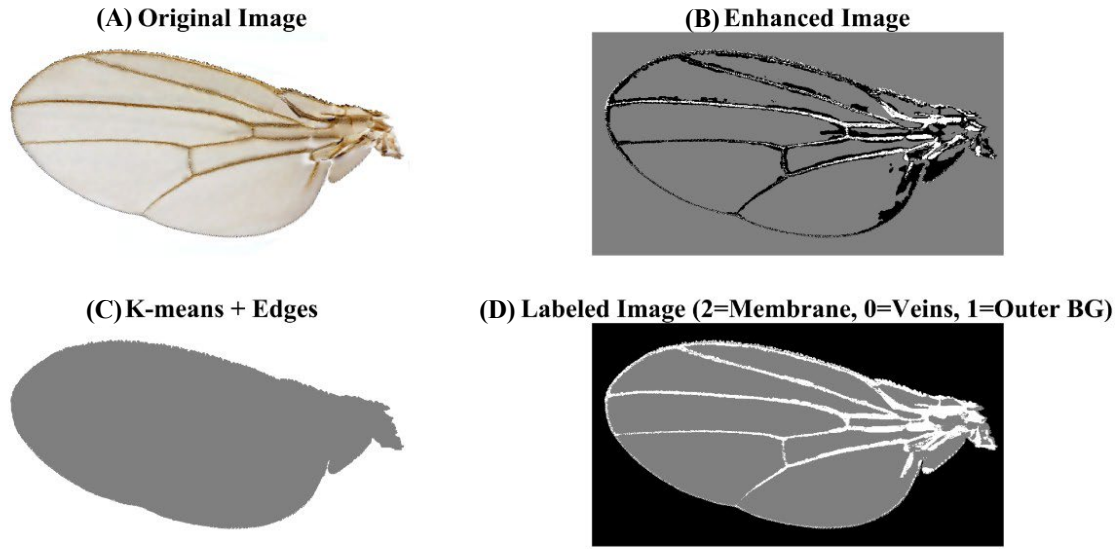

**Fig. S1. Membrane and vein pixel identification workflow.** (A) RGB image of a *D. melanogaster* wing used as the input for segmentation, obtained from Flickr (Jones, 2013). (B) Contrast-enhanced grayscale image obtained from grayscale conversion and application of Contrast Limited Adaptive Histogram Equalization (CLAHE), improving the visibility of veins and wing boundaries. (C) Preliminary segmentation combining k-means clustering in RGB color space with edge detection to isolate the wing region from the background. (D) Final labeled image after morphological refinement: wing membrane (label 2), vein structures (label 0), and outer background (label 1).

### S1.2 Membrane thickness mapping

In this study, membrane thickness is estimated using an inverse colourimetry approach, in which a thin-film optical model is fitted to the reflectance spectra inferred from RGB WIP images (Li et al., 2022). We obtained The WIP image of a *D. melanogaster* wing from the dataset reported in De Lisle et al. (2025). To ensure reliable thickness estimation, we first apply a pixel-level filtering procedure to the WIP image to exclude regions with low signal quality. Pixels are classified as unreliable based on color properties in the HSV (Hue, Saturation, Value) color space, enabling the removal of dark pixels, desaturated regions, and ambiguous color responses associated with noise, reflections, or non-interference artefacts. The resulting binary mask is further refined using median filtering to suppress isolated misclassifications. Only pixels passing this color-based filtering are retained for subsequent analysis ( Fig. S2A). The filtered WIP image is spatially registered to the anatomical reference RGB image to ensure pixel-wise correspondence between thickness estimates and wing geometry (Fig. S2B), using manually selected control points placed at vein intersections and along wing boundaries, distributed symmetrically in both chordwise and spanwise directions. A third-order polynomial transformation is then applied to warp the WIP image onto the anatomical reference frame.

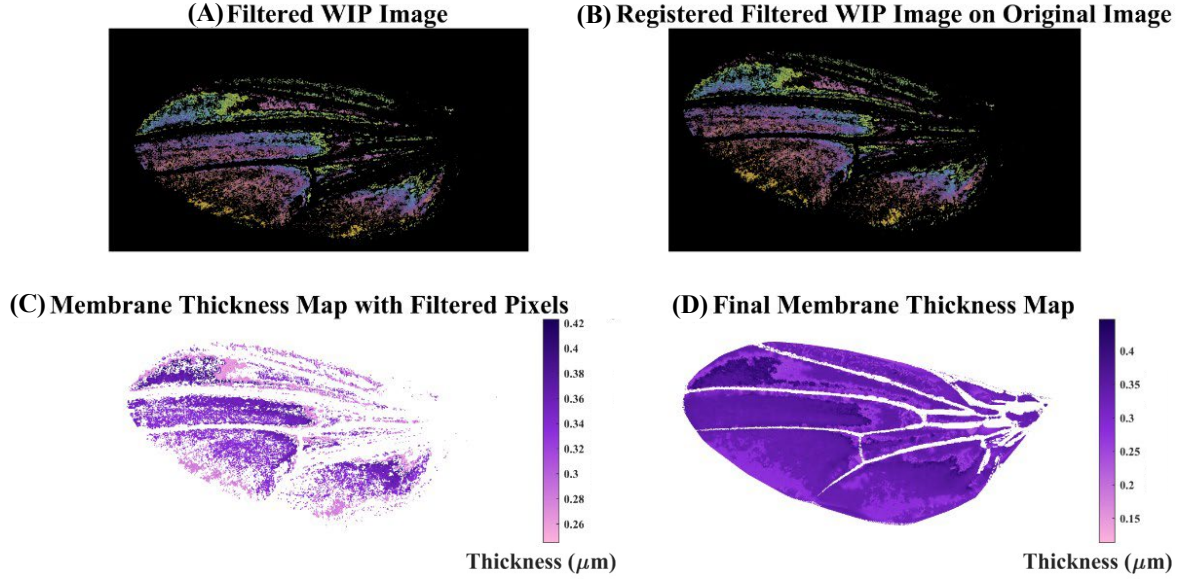

**Fig. S2. Workflow for Membrane thickness mapping from WIP imagery.** (A) Filtered WIP image after HSV-based pixel quality filtering and median refinement. (B) Spatial registration of the filtered WIP image onto the anatomical reference wing image using manually selected control points and a third-order polynomial transformation. (C) Sparse membrane thickness map obtained by fitting a thin-film interference model to the reconstructed reflectance spectra of the filtered pixels. (D) Final membrane thickness map after residual-error filtering, cubic interpolation of missing values, and masking with the segmented wing domain.

For each valid pixel in the registered WIP image, the recorded RGB triplet is normalised and used to infer the underlying reflectance spectrum. This is achieved by modelling the camera's spectral sensitivities as Gaussian response functions for the red, green, and blue channels. A non-negative least-squares optimisation is then employed to estimate the most likely reflectance intensity at each wavelength. The inferred reflectance spectrum is evaluated over the visible wavelength range of 450–650 nm, corresponding to the effective transmission range of standard digital camera sensors equipped with UV–IR cut-off filters. The reconstructed reflectance spectrum at each pixel is fitted to a thin-film interference model, which is expressed as (Li et al., 2022):

$$F(\gamma, d) = \frac{4R_s \sin^2 \alpha}{(1 - R_s)^2 + 4R_s \sin^2 \alpha}, \quad \alpha = \frac{2\pi d \sqrt{n^2 - \sin^2 \theta}}{\gamma}, \quad (1)$$

where $n = 1.57$ is the refractive index of chitin; $d$ is the membrane thickness; $\theta$ is the incidence angle, which, for the experimental conditions of De Lisle et al. (2025), is normal ($\theta = 0^\circ$); $\gamma$ is the wavelength; and $R_s$ is the reflection coefficient for s-polarised light:

$$R_s = \left( \frac{\cos \theta - \sqrt{n^2 - \sin^2 \theta}}{\cos \theta + \sqrt{n^2 - \sin^2 \theta}} \right)^2. \quad (2)$$

Thickness values are estimated by fitting the model to the inferred reflectance spectrum using constrained non-linear least-squares optimization. In some pixels, noisy, desaturated, or spectrally ambiguous colour responses resulted in poor agreement between the thin-film model and the inferred reflectance spectrum. These cases are identified using the residual fitting error, quantified by the sum of squared residuals. This filtering step complements the initial HSV-based pixel selection and reduces the influence of non-physical or non-convergent thickness estimates (Fig. S2C). Following quality control, the resulting thickness values forms a sparse map over the wing membrane, and missing values are interpolated using cubic interpolation, which is chosen due to the continuous and smoothly varying nature of thin-film membrane thickness. Cubic interpolation provides second-order smoothness, avoids artificial discontinuities, and more accurately preserves gradual thickness gradients than linear methods, particularly in regions corresponding to interference fringes. Finally, pixels outside the wing domain are removed using a binary wing mask derived in the previous section. The final membrane thickness map, restricted to the wing domain is shown in Fig. S2D.

#### **S1.3 Characterization of the mass density of the membrane and veins**

##### ***Membrane density considerations:***

In many dipteran, hymenopteran, and lepidopteran insects, the wing membrane does not consist of a single homogeneous layer. Instead, it is formed by two extremely thin and transparent cuticular laminae, referred to as the dorsal and ventral veils, which become closely apposed during development and collectively give the wing the appearance of a single translucent surface (Sun and Bhushan, 2012; Zhao et al., 2014). So, the membrane's density is expected to be close to that of the constituent dry chitin-protein composite materials, *i.e.*, in the range of approximately 1.3–1.5 g cm<sup>-3</sup>. Although the wing membrane requires only very low hydration (Pass, 2018), water uptake significantly influences its mechanical properties (Enders et al., 2004; Klocke and Schmitz, 2011a; Klocke and Schmitz, 2011b), including a reduction in the bulk density to about 85-90% of the dry cuticle density, yielding a hydrated density in the range of 1.1-1.25 g cm<sup>-3</sup>. In the present study, the membrane volumetric density is assumed to be spatially uniform and is assigned a value of 1.2 g cm<sup>-3</sup>, consistent with previously reported measurements (Klocke and Schmitz, 2011b).

##### ***Vein structural composition:***

Insect wing veins are not merely empty cuticular tubes; they are highly integrated, multilayered conduits that combine structural, sensory, and transport functions. The longitudinal veins form the main scaffolding of the wing, each consisting of a thin chitin-protein cuticular shell that surrounds a central lumen filled with hemolymph, tracheal tubes, nerves and an outer epidermal layer (Pass, 2018). Resilin-rich elastic pads are also present in the internal layers of the vein cuticle, requiring continuous hydration to retain their elasticity (Pass, 2018). The substances that flow through these veins are essential for wing function. Hemolymph circulates through the venous network, delivering water, nutrients, hormones and dissolved oxygen to the living tissues of the wing while removing metabolic waste (Pass, 2018). In addition, a network of tracheae runs within the veins, allowing gaseous exchange between the ambient air and the hemolymph (Sugiyama et al., 2024). Water supplied by the hemolymph also hydrates the cuticle of the veins, preserving its elasticity and preventing desiccation-induced brittleness (Pass, 2018). These structural and fluid-transport components together dictate the mechanical properties of the wing veins. In particular, the bulk density of veins is expected to be lower than that of solid cuticle owing to the presence of low-density haemolymph and, in many species, air-filled tracheae within the hollow vein lumen. Consequently, the vein's effective density is a composite value that reflects both its stiff cuticular walls and the lighter internal fluids.

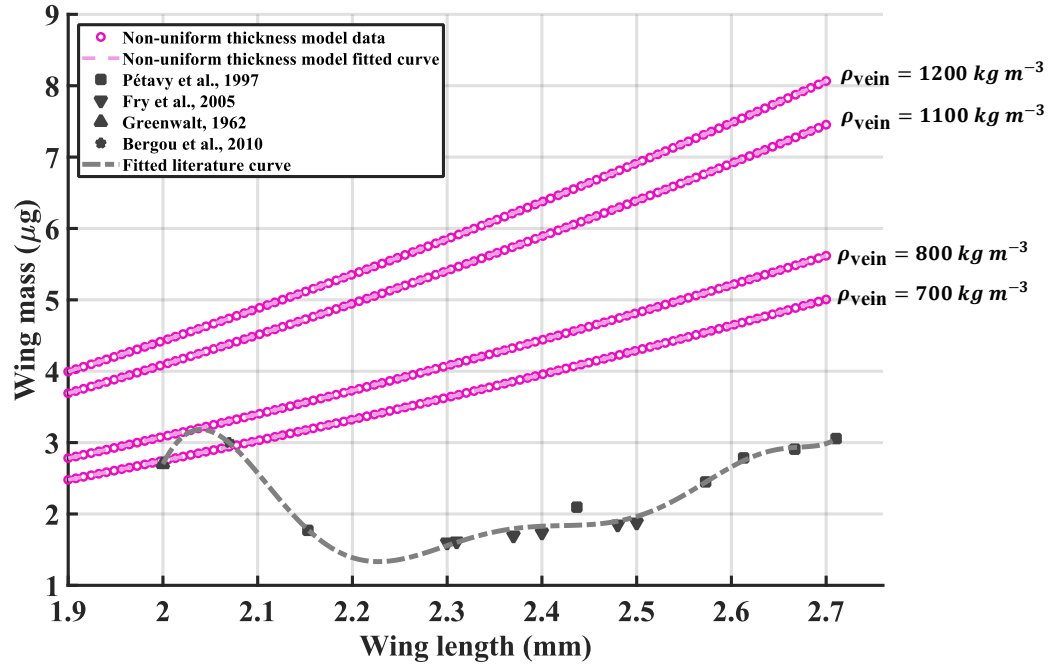

**Fig. S3. Calibration process for Determining the effective wing vein density.** Model-predicted total wing mass as a function of wingspan length for several assumed vein density values. The model results are compared with experimentally reported wing mass measurements for *D. melanogaster* from the literature (Bergou et al., 2010; Fry et al., 2005; Greenwalt, 1962; Lehmann and Dickinson, 1997; Pétavy et al., 1997).

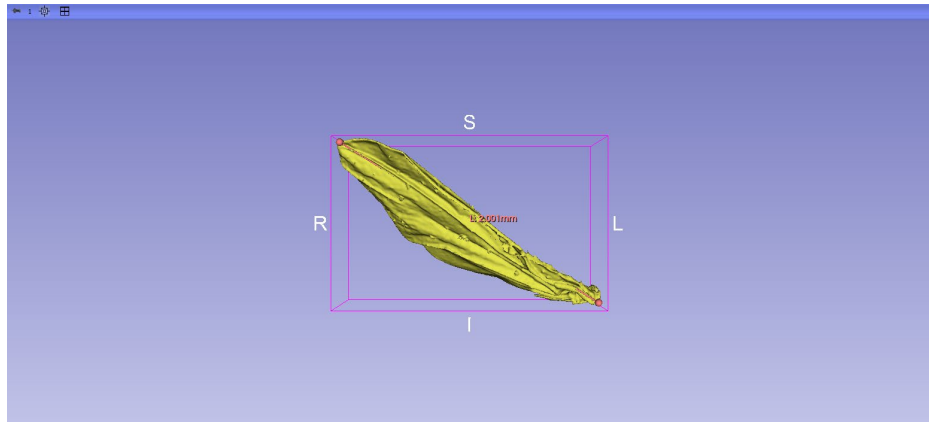

**Fig. S4. Measurement of the wingspan length from the $\mu\text{CT}$-reconstructed three-dimensional wing model.** Three-dimensional geometry of the *D. melanogaster* wing is reconstructed from the micro-CT volume, reported in Maisano (2024), and exported as an STL surface model. The span length of the wing is measured directly within the 3D Slicer environment along the spanwise direction, yielding a value of approximately 2.0 mm. This measurement is used to identify the physically relevant intersection point between the model-predicted wing mass and the fitted literature trend, ensuring consistency between the $\mu\text{CT}$ derived morphology and the optical wing image used in the present analysis.

##### ***Calibration of effective vein density:***

To determine an appropriate value for the effective density of the wing vein structure corresponding to the reference wing analysed through its optical image, we implement the following workflow. A range of plausible vein density values is first selected, and for each assumed density the total wing mass is calculated over a biologically relevant span length range of 1.9-2.7 mm. The resulting total wing masses are then plotted as a function of span length and compared with experimental data reported in the literature for *D. melanogaster* (Bergou et al., 2010; Fry et al., 2005; Greenewalt, 1962; Lehmann and Dickinson, 1997; Pétavy et al., 1997). Fitted curves are generated for both the model predictions and the scattered literature measurements. As shown in Fig. S3, the model curve corresponding to a vein density of $700 \text{ kg m}^{-3}$ exhibits the closest agreement with the fitted literature trend. Two intersections between the model and literature curves are identified. The first occurs at a span length of 2.0018 mm, corresponding to a total wing mass of $2.747 \mu\text{g}$, while the second occurs at a span length of 2.0745 mm. The first intersection is selected as the physically relevant solution. This choice is guided by independent morphological measurements derived from micro-CT data—the three-dimensional wing model reconstructed from the micro-CT volume (Maisano, 2024) and analysed in 3D Slicer—where direct measurement of the wingspan yields a value of approximately 2.0 mm, as shown in Fig. S4. Selecting this span length ensures consistency between the wing morphology represented in the micro-CT dataset and the biological wing depicted in the optical image. The corresponding wing mass at this intersection point is therefore used to compute the effective vein density. This value, which is $700 \text{ kg m}^{-3}$, is subsequently adopted as the representative vein density for all further analyses.

##### **S1.4 Spanwise mass and inertial distribution in lower-fidelity models**

Two models with increasing simplification are implemented and compared with the developed pixel-based non-uniform thickness distribution model. All models compute spanwise quantities by integrating pixel-wise contributions along columns of the optical wing image. The comparison assesses the influence of spatial variations in thickness, density, and geometry on wing inertial properties and the effects of simplifying assumptions commonly employed in reduced-order wing models on flight motor predictions (Fig. 7 in the main text). The optical wing image is discretised into spanwise columns ($i = 1, \dots, N_x$). Within each column, pixel contributions are summed over the chordwise direction ($j = 1, \dots, N_y$). Fractional pixel occupancy, $f_{j,i}$, accounts for boundary corrections, either in the vein mask ($f_{j,i}^v$) or membrane mask ($f_{j,i}^m$), $r_{j,i}$ denotes the distance from the wing root, and $t_{j,i}^v$ and $t_{j,i}^m$ represent the local vein and membrane thicknesses, respectively. $N_p$ show the total number of pixels within the whole wing, the vein mask and the membrane mask, respectively.

#### ***Uniform plate model***

To isolate the effect of thickness variation, the wing is modelled as a uniform plate with mean thickness,

$$t_{\text{eff}} = \frac{\sum_{j,i} t_{j,i}}{N_p}, \quad (3)$$

and constant surface density,

$$\sigma = \rho t_{\text{eff}}, \quad \rho = \rho_{\text{cuticle}} = 1200 \text{ kg m}^{-3}. \quad (4)$$

Spanwise mass and inertia are then obtained from:

$$\begin{aligned} m_{i,k} &= \sigma A_p \sum_{(j,i) \in \Omega_k} f_{j,i}^k, \\ I_{i,k} &= \sigma A_p \sum_{(j,i) \in \Omega_k} (f_{j,i}^k r_{j,i}^2), \end{aligned} \quad (5)$$

where $\Omega_k | k \in \{\text{wing, vein, membrane}\}$ denotes the selected region of interest.

#### ***Uniform rectangular prism model***

Finally, the wing is approximated as a uniform rectangle with constant mean thickness and density. Spanwise mass and inertia are computed using the same relations expressed in Eqn 5. This model neglects all spatial variation in thickness, material composition, and planform geometry, serving as a lower-bound reference for model fidelity.

### **S2 Effective mass and inertia of thoracic deformation**

#### **S2.1 Thoracic geometry reconstruction**

A micro-CT dataset for *D. melanogaster* is obtained from the digital repository MorphoSource (Maisano, 2024). These data are processed in 3D Slicer to reconstruct detailed three-dimensional thoracic structures. The initial output is a point cloud representing both internal and external surfaces, which is needed to be converted into a continuous surface mesh using MeshMixer through the following steps: **(i)** Surface reconstruction by triangulation of discrete points. **(ii)** Noise reduction and smoothing to remove imaging artefacts while preserving anatomical features. And **(iii)** Mesh repair to ensure watertight geometry by filling holes and correcting non-manifold edges. The processed mesh is imported into Blender, where thoracic components are aligned, assembled, and scaled; and geometric inconsistencies such as intersecting surfaces are corrected.

This workflow, (3D Slicer – MeshMixer – Blender), ensures that the created thorax model, as shown in Fig. 3 of the main text, preserves anatomical realism while being suitable for mechanical and vibrational analysis. To account for major sclerites in the thorax model, a detailed 3D model is created following the previously described CAD workflow, and the complete geometry is exported as an STL file from Blender. The thorax is modelled as a solid 3D domain, while the sclerites are represented using shell elements. These shells are connected to the solid thorax domain via a thin-to-solid coupling interface, ensuring proper transmission of forces and displacements between the components.

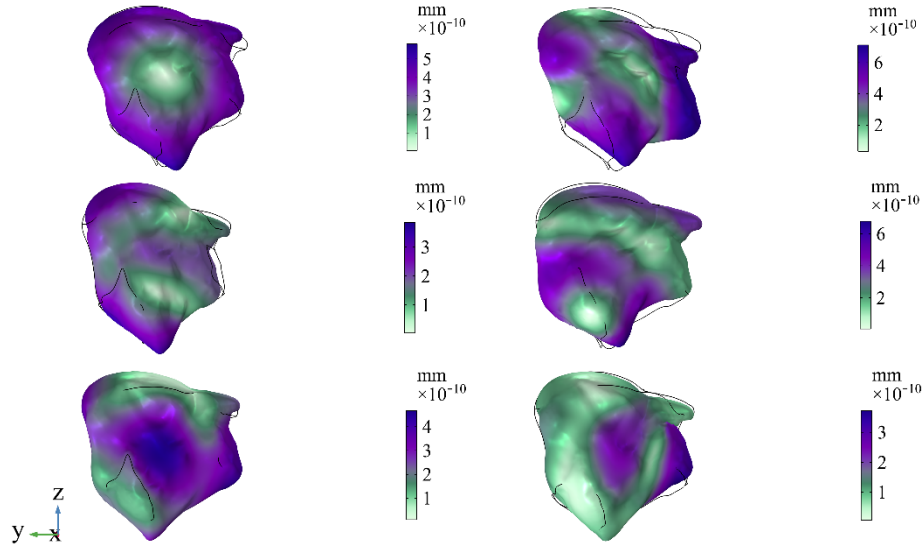

**Fig. S5. FEA modal results.** Six different mode shapes obtained through COMSOL Multiphysics analysis for the sample *D. melanogaster* thorax, arranged left to right, top to bottom in numerical order.

### S2.2 Modal analysis procedure in COMSOL

The geometry, comprising both the thorax and associated sclerites, is imported into COMSOL for analysis, where a Multiphysics framework is employed to account for the composite nature of the structure. For the simulation, symmetry along the sagittal plane is assumed, and thus only half of the thorax is modelled. To investigate the effective mass or the effective rotational inertia of the thorax resulting from its vibration, we must estimate the thoracic deformation patterns. These representative patterns are obtained through modal analysis, in which the eigenfrequency problem is solved numerically using the finite-element method. The procedure consists of discretizing the geometry into finite elements, assigning the appropriate material properties, including Young's modulus, Poisson's ratio, and density, and subsequently solving the eigenfrequency problem to obtain the natural frequencies, displacement fields, and corresponding mode shapes.

The resulting modes are mutually orthogonal and represent the distinct deformation patterns through which the thoracic structure can vibrate, with each mode characterized by a specific natural frequency and spatial displacement field. Modal analysis of the *D. melanogaster* thorax identifies a total of six mode shapes, whose corresponding deformation patterns are illustrated in Fig. S5. Given the presence of both solid and shell elements with differing characteristics, the degrees of freedom (DOFs) are used as the metric to evaluate mesh dependency and convergence of the results. The mesh independence study for the *D. melanogaster* thorax model is shown in Fig. S6, where the computed natural frequency is plotted as a function of the total degrees of freedom (DOFs). As the mesh is refined, the frequency converges toward a stable value near 3.5 kHz. Beyond approximately $5 \times 10^5$ DOFs, further mesh refinement produces negligible changes in the computed frequency. This convergence confirms that the solution is mesh-independent.

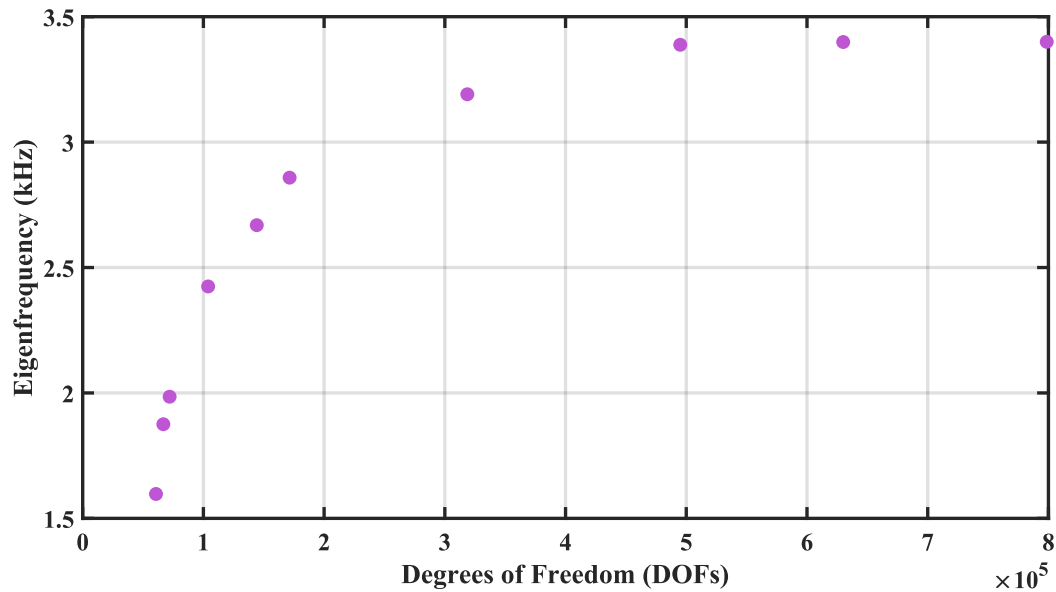

**Fig. S6. Mesh independence study for the *D. melanogaster* thorax model.** Computed natural frequency of the first vibration mode as a function of the total degrees of freedom (DOFs) used in the finite element discretisation.

#### S3 Meta-analysed unilateral wing clipping data

A meta-analysis conducted using experimental datasets from multiple dipteran species to evaluate the response of wingbeat frequency to unilateral wing damage, and to compare these responses with dynamic model predictions. In Salem et al. (2022), raw frequency measurements for rigidly tethered *D. melanogaster* with damaged wings are reported; however, intact-state frequencies are not provided for individual specimens. Instead, a single average intact wingbeat frequency derived from 16 specimens is presented. Consequently, frequency ratios between damaged and intact states are calculated using this reported mean value as the reference. Additional data are obtained from Muijres et al. (2017), in which supplementary dataset S1 (rsfs20160103supp2.mat) contains wing morphology and flight kinematics for free-flying *D. hydei* subjected to unilateral wing damage. Within the wing\_kinematics structure, the variables n\_down\_start\_L and n\_down\_start\_R correspond to the frame indices marking the onset of successive downstroke cycles for the left and right wings, respectively. Given that the recordings were acquired at a known frame rate, the time interval between consecutive downstroke initiations was computed, allowing the wingbeat period and corresponding frequency to be determined for each wing. Accordingly, for each specimen, the relevant damaged wing (identified through the clip\_side variable) frequency, together with the corresponding wing area ratio (residual wing area after damage divided by the intact wing area) of the respective cut experiment, either spanwise or chordwise clip cases, were identified. Since we required the normalised frequencies of the damaged states relative to their respective intact states for comparison with the proposed PEA models, we sought the specimen-specific intact frequencies. However, no individual intact frequency data are reported in the kinematics dataset.

Instead, the mean ± sd of the wingbeat frequency calculated from 1603 steady wingbeats across 83 undamaged flies is provided (denoted as $f_{\text{normal}}$). So, this reported mean value is adopted as the reference frequency for normalisation.

In Deora et al. (2021), wingbeat frequency of single-wing clipped *Hermetia illucens* flies is reported as a function of normalized wing length. To enable comparison of these data, we converted these normalized wing lengths to normalized wing areas, based on the normalized wing area as a quartic function of normalized wing lengths derived from the optical reference wing image in the present study. This relationship was assumed to provide a reasonable approximation of the geometric scaling of *Hermetia illucens* wings. Among the morphological data reported in Table 1 of Hao et al. (2025) for *Eristalis tenax*, the wing area is not explicitly stated. The wing area is therefore approximated as the product of wing span length and mean chord length. Using the reported aspect ratio, defined as the ratio of wing span length to mean chord length, the corresponding wing area is estimated. In Sotavalta (1953), the mutilated wing length ratio in midges (*Chironomus* s.lat.) is reported, whereas no information is provided regarding the remaining wing area ratio after transverse cuts. The area of the truncated wing relative to the intact wing is therefore approximated by analysing the *Chironomus* wing morphology using *ImageJ*.

### S4 Hybrid elastic actuation (HEA) with wing root flexion

#### S4.1 Dynamics of the HEA model

The equations of motion of the hybrid system introduced in the main text are

$$\begin{aligned} M\ddot{x} + D\dot{x} &= k_s(u(t) - x(t)), \\ F(t) - k_p u(t) &= k_s(u(t) - x(t)), \end{aligned} \quad (6)$$

or, equivalently:

$$M\ddot{x} + D\dot{x} = F(t) - k_p u(t). \quad (7)$$

In addition to parameters from the PEA model, this HEA model contains a series elasticity $k_s$, defining a natural frequency ($\Omega_{0,s}$) and damping ratio ($\xi_s$):

$$\Omega_{0,s} = \sqrt{\frac{k_s}{M}}, \quad \xi_s = \frac{D}{2\sqrt{Mk_s}}. \quad (8)$$

To quantify the relative strength of the series elasticity with respect to the parallel elasticity, the non-dimensional parameter $\alpha$ is introduced:

$$\alpha = \frac{\Omega_{0,s}}{\Omega_{0,p}} = \sqrt{\frac{k_s}{k_p}}. \quad (9)$$

So, with the definition of $\alpha$, the relation between damping ratios is:

$$\xi_p = \alpha \xi_s. \quad (10)$$

Considering the canonical representation of the equations of motion, from second relation in Eqn.6, $u(t)$ could be represented as a function of $f(t)$ and $x(t)$:

$$u(t) = \frac{f(t) + \Omega_{0,s}^2 x(t)}{\Omega_{0,s}^2 + \Omega_{0,p}^2}. \quad (11)$$

which, upon substitution into Eqn 7. gives:

$$\ddot{x} + 2\xi_p \Omega_{0,p} \dot{x} = f(t) - \Omega_{0,p}^2 \left( \frac{f(t) + \Omega_{0,s}^2 x(t)}{\Omega_{0,s}^2 + \Omega_{0,p}^2} \right), \quad (12)$$

and, Eqn 6. is formulated as:

$$\ddot{x} + 2\xi_p \Omega_{0,p} \dot{x} = \Omega_{0,s}^2 (u(t) - x(t)). \quad (13)$$

To evaluate the global resonant behaviour, the system is analysed in the frequency domain. The time-varying quantities are expressed in complex (phasor) form as:

$$x(t) = X \cos(\Omega t + \psi_x) = \Re\{\hat{x} e^{i\Omega t}\} = |\hat{x}| \Re\{\exp(i\Omega t + i\psi_x)\}, \quad \psi_x = \tan^{-1} \left( \frac{\Im\{\hat{x}\}}{\Re\{\hat{x}\}} \right),$$

with:

$$\begin{aligned} u(t) &= \Re\{\hat{u} e^{i\Omega t}\}, \\ f(t) &= \Re\{\hat{f} e^{i\Omega t}\}, \\ \dot{u}(t) &= \Re\{\hat{u} i\Omega e^{i\Omega t}\}. \end{aligned} \quad (14)$$

Here, $\hat{x}$, $\hat{u}$ and $\hat{f}$ denote complex amplitudes containing both real-valued magnitude and phase angle, and $\Omega$ is the excitation frequency. Without loss of generality, $\hat{f}$ is taken as real ($\hat{f} \in \mathbb{R}$).

Substituting the phasors into Eqns 12 and 13, allows us to define two transfer functions between input force and output displacement and between input and output displacements respectively:

$$\frac{\hat{x}}{\hat{f}} = \frac{\frac{\Omega_{0,s}^2}{\Omega_{0,s}^2 + \Omega_{0,p}^2}}{-\Omega^2 + 2\xi_p \Omega_{0,p} i\Omega + \frac{\Omega_{0,p}^2 \Omega_{0,s}^2}{\Omega_{0,s}^2 + \Omega_{0,p}^2}}, \quad (15)$$

$$\frac{\hat{x}}{\hat{u}} = \frac{\Omega_{0,s}^2}{-\Omega^2 + 2\xi_p \Omega_{0,p} i\Omega + \Omega_{0,s}^2}.$$

These two transfer functions enable characterization of the mechanical power consumption of the flight motor system and identification of the frequencies at which power expenditure is minimised. These frequencies are referred to as the global resonant frequencies (Ma and Zhang, 2014; Pons and Beatus, 2022), which correspond to the effective resonant state in the hybrid flight motor self-oscillation (Pons, 2023).

Defining the instantaneous mechanical power requirement of the system in the canonical form $p = f\dot{u}$, the result is composed of a constant (DC) component and a second-harmonic component, and it cannot be represented by a single phasor (Findeisen and Popp, 2002). Using the phasor representations introduced in Eqn 14, the power could be represented as below, with the constant term and the oscillatory term, as the first and second ones respectively:

$$p(t) = p_0 + \Re\{\hat{p}e^{2i\Omega t}\}. \quad (16)$$

Defining the canonical parameters $f$ and $\dot{u}$ with their complex conjugates $(.)^\dagger$ leads to:

$$\begin{aligned} f(t) &= \frac{1}{2}(\hat{f}e^{i\Omega t} + \hat{f}^\dagger e^{-i\Omega t}), \\ \dot{u}(t) &= \frac{1}{2}(\hat{v}e^{i\Omega t} + \hat{v}^\dagger e^{-i\Omega t}). \end{aligned} \quad (17)$$

Substitution into the power expression yields:

$$p(t) = \underbrace{\frac{1}{4}(\hat{v}^\dagger \hat{f} + \hat{v} \hat{f}^\dagger)}_{p_0} + \underbrace{\frac{1}{4}(\hat{v} \hat{f} e^{2i\Omega t} + (\hat{v} \hat{f} e^{2i\Omega t})^\dagger)}_{\Re\{\hat{p}e^{2i\Omega t}\}}. \quad (18)$$

The oscillatory power term could be converted into explicit trigonometric form as:

$$\Re\{\hat{p}e^{2i\Omega t}\} = |\hat{p}| \cos(2\Omega t + \psi_p) = |\hat{p}_1| \sin(2\Omega t) + |\hat{p}_2| \cos(2\Omega t), \quad (19)$$

with amplitude:

$$|\hat{p}| = \sqrt{|\hat{p}_1|^2 + |\hat{p}_2|^2}. \quad (20)$$

To express the power components $p_0$ and $|\hat{p}|$ in terms of the input muscular force amplitude ($\hat{f} \in \mathbb{R}$), the chain of transfer functions defined in Eqn 15. is employed:

$$\frac{\hat{u}}{\hat{f}} = \left(\frac{\hat{x}}{\hat{u}}\right)^{-1} \left(\frac{\hat{x}}{\hat{f}}\right). \quad (21)$$

This substitution leads to explicit analytical expressions for the normalised constant power component and oscillatory amplitude:

$$\begin{aligned} \frac{p_0}{\hat{f}^2} &= \frac{\xi_p \Omega_{0,p} \Omega^2 \Omega_{0,s}^4}{(\Omega_{0,p}^2 \Omega_{0,s}^2 - (\Omega_{0,s}^2 + \Omega_{0,p}^2) \Omega^2)^2 + (2\xi_p \Omega_{0,p} \Omega (\Omega_{0,s}^2 + \Omega_{0,p}^2))^2}, \\ \frac{|\hat{p}|}{\hat{f}^2} &= \frac{\Omega}{2} \sqrt{\frac{(\Omega_{0,s}^2 - \Omega^2)^2 + (2\xi_p \Omega_{0,p} \Omega)^2}{(\Omega_{0,p}^2 \Omega_{0,s}^2 - (\Omega_{0,s}^2 + \Omega_{0,p}^2) \Omega^2)^2 + (2\xi_p \Omega_{0,p} \Omega (\Omega_{0,s}^2 + \Omega_{0,p}^2))^2}}. \end{aligned} \quad (22)$$

Energy (or, global) resonance (Pons and Beatus, 2022; Zhang et al., 2018) is achieved when the overall power balance of the system is structured such that no net negative work appears in its total energy requirement:

$$p(t) \geq 0, \forall t. \quad (23)$$

The satisfaction of this condition at a particular frequency renders the system optimal in terms of energetic demand concept. Denoting the energy-resonant frequency of our hybrid model as $\Omega_g$, condition (23) could be translated into:

$$\Omega_g = \{\Omega: p_0 \geq |\hat{p}|\}. \quad (24)$$

Via relations 22, $p_0 > |\hat{p}|$ is never satisfied by any real $\Omega$. Thus, energy resonance can occur only under equality conditions:

$$\Omega_g = \{\Omega: p_0 = |\hat{p}|\}. \quad (25)$$

Solving this equation yields two analytical solutions in terms of the series parameters as:

$$\Omega_g^\pm = \Omega_{0,s} \mathcal{G}^\pm(\alpha, \xi_s),$$

$$\mathcal{G}^\pm(\alpha, \xi_s) = \frac{\sqrt{\pm \sqrt{A_1(\alpha, \xi_s) B_1(\alpha, \xi_s)} + C_1(\alpha, \xi_s)}}{\sqrt{2(\alpha^2 + 1)}}, \quad (26)$$

Where

$$\begin{aligned} A_1(\alpha, \xi_s) &= \alpha^2(1 - 2\xi_s)^2 + 4\xi_s(\xi_s - 1), \\ B_1(\alpha, \xi_s) &= \alpha^2(1 + 2\xi_s)^2 + 4\xi_s(\xi_s + 1), \\ C_1(\alpha, \xi_s) &= 2 + \alpha^2 - 4\xi_s^2(1 + \alpha^2). \end{aligned}$$

##### S4.2 Numerical validation of analytical self-oscillation frequencies

The two self-oscillation frequency solutions $\Omega_g^+$, $\Omega_g^-$ correspond to SEA-like (wing hinge-dominated) resonance and PEA-like (thorax-dominated) resonance behaviour respectively. These analytical solutions of the frequencies are needed to be validated. First, we define two complementary metrics of mechanical power transfer efficiency that quantify how effectively muscular input power is converted into useful net power throughput by the flight motor. The absolute mechanical power transfer efficiency, $\eta_{abs}$ is defined as the ratio of the net mechanical work performed over a cycle to the total absolute power expenditure:

$$\eta_{abs} = \frac{\int_0^T p(t) dt}{\int_0^T |p(t)| dt} \quad (27)$$

This metric represents the efficiency of an actuator without energy regeneration, for which both positive and negative work contributions incur energetic cost. In contrast, the positive-only mechanical power transfer efficiency, $\eta_{pos}$, is defined as:

$$\eta_{\text{pos}} = \frac{\int_0^T p(t)dt}{\int_{\{t \in [0,T]: p(t) > 0\}} p(t)dt}. \quad (28)$$

This formulation accounts only for positive power input in the denominator and thus corresponds to a system capable of dissipating negative work without recovering it. In both cases, the numerator represents the net mechanical power transmitted through the system, while the denominator represents a measure of muscular energetic cost. The efficiencies attain their maximal value of unity when the instantaneous power remains non-negative throughout the cycle, that is, under the condition of energy resonance, where negative work absorption is eliminated. Together, these definitions provide a rigorous framework for evaluating energetic optimality in oscillatory flight motor dynamics. Numerical evaluation demonstrates that the poles of the power transfer efficiencies coincide with the analytically derived energy resonant frequencies, thereby validating the analytical formulation (Fig. S7).

#### S4.3 Scaling of linear damping ratio with wing damage

Considering the linear damping ratio $\xi_s$, defined as Eqn 8, we first determine the equivalent viscous damping parameter in 1DOF rotational frame to account for the wings inertia directly. The rotational version of the Eqn 6. is:

$$I\ddot{\theta} + D_\theta \dot{\theta} = k_{s,\theta}(\phi(t) - \theta(t)), \quad (29)$$

where the torque applied to the wing hinge arises from the rotational displacement of input angle $\phi(t)$ relative to the wing hinge angle $\theta(t)$. Rotational viscous damping ratio is defined as:

$$\xi_s = \frac{D_\theta}{2\sqrt{Ik_{s,\theta}}}. \quad (30)$$

We consider the wing model as a rectangle for simplification as study (Salem et al., 2022) did. In Eqn 29, the term $D_\theta \dot{\theta}$ refers to the aerodynamic drag torque which is equal to the normal drag force multiplied by the horizontal arm distance to the wing hinge (wingspan length $\ell$):

$$\tau_D = F_D \ell = \frac{1}{2} \rho C_D A u^2 \ell = \frac{1}{2} \rho C_D \ell h (\ell \dot{\theta})^2 \ell, \quad (31)$$

$$\tau_D = \frac{1}{2} \rho C_D h \ell^4 \dot{\theta}^2,$$

where $\rho$ denotes air density, $C_D$ a drag coefficient, $A$ an effective aerodynamic area and $u$ aerodynamic velocity. So, the damping parameter $D_\theta$ yields to:

$$D_\theta = \frac{1}{2} \rho C_D h \ell^4 \dot{\theta} = \frac{1}{2} \rho C_D h \ell^4 \Omega. \quad (32)$$

For the simplified rectangular wing model, the wing inertia is:

$$I = \frac{\rho h \ell^3 w}{3}. \quad (33)$$

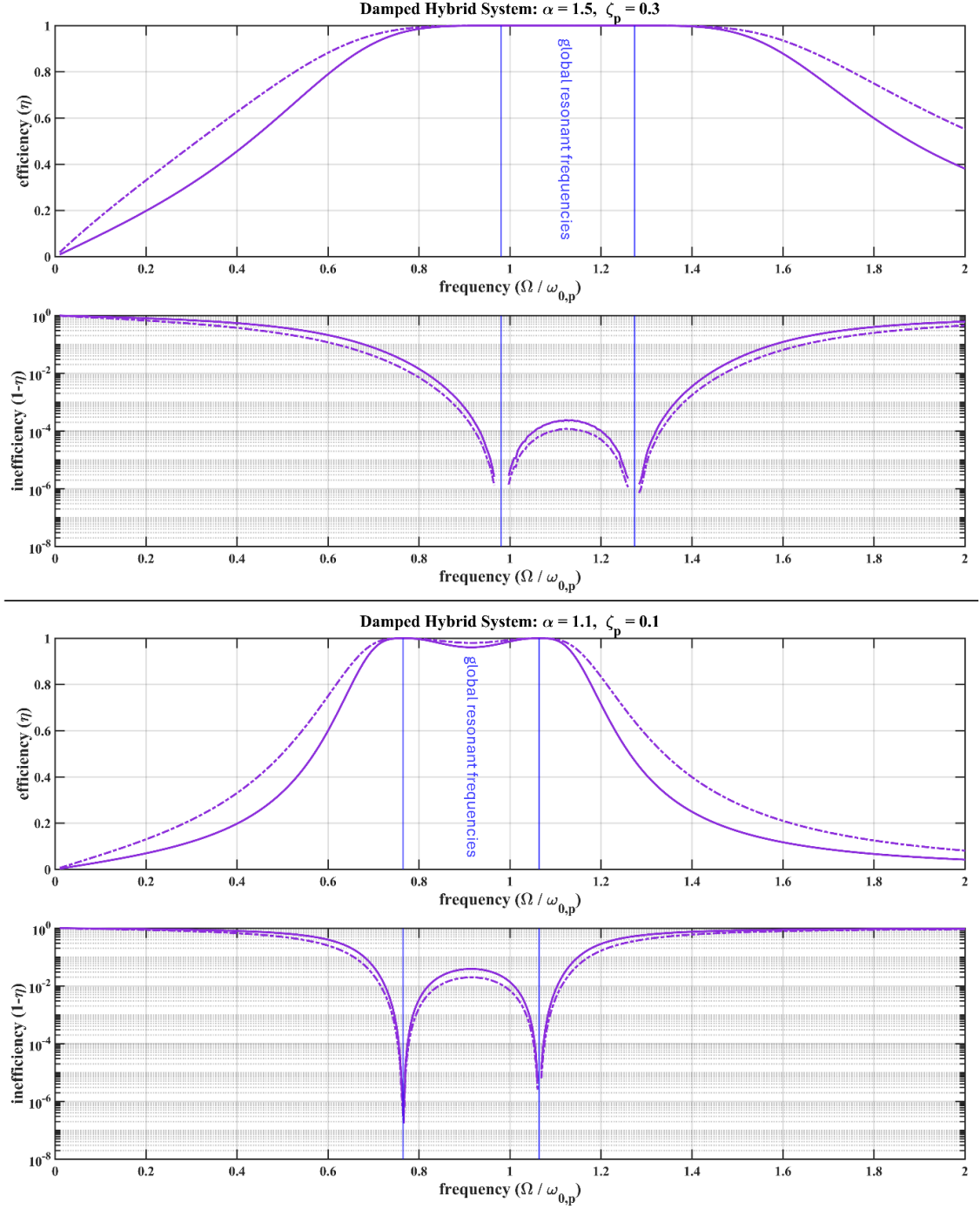

**Fig. S7. Numerical validation of analytically predicted global resonant frequencies in the hybrid model, using mechanical power transfer efficiency.** The absolute efficiency $\eta_{abs}$ and positive-only efficiency $\eta_{pos}$ are plotted as functions of the normalized excitation frequency $\Omega / \omega_{0,p}$ for two representative HEA models. The upper panels show the efficiencies, while the corresponding lower panels present their reciprocal forms (inefficiency) to highlight the singular behaviour near resonance.

Therefore, the rotational viscous damping ratio could be rewritten as:

$$\xi_s = \frac{\frac{1}{2} \rho C_D h \ell^4 \Omega}{2 \sqrt{\frac{\rho h \ell^3 w}{3} k_{s,\theta}}} = \frac{\sqrt{3}}{4} C_D \frac{\sqrt{\rho h}}{\sqrt{w k_{s,\theta}}} \ell^{\frac{5}{2}} \Omega. \quad (34)$$

Thus:

$$\xi_s \propto \ell^{\frac{5}{2}} \Omega, \quad \frac{\xi_{s,\text{dam}}}{\xi_{s,\text{int}}} \propto \left( \frac{\ell_{\text{dam}}}{\ell_{\text{int}}} \right)^{\frac{5}{2}} \frac{\Omega_{\text{dam}}}{\Omega_{\text{int}}} \propto \left( \frac{I_{\text{dam}}}{I_{\text{int}}} \right)^{\frac{5}{6}} \frac{\Omega_2}{\Omega_1}. \quad (35)$$

Chordwise trimming of the wings from tip to the distal part, leaves the rectangular model wing chord $h$ and wing width $w$ constant, while changes the span length $\ell$ and flapping frequency $\Omega$. Based on the mutilation experiments conducted in (Sotavalta, 1952), among dipteran species the wing stroke frequency has a relation with the total moment of inertia of the wings with the mean value above exponent with:

$$\frac{\Omega_2}{\Omega_1} \propto \left( \frac{I_2}{I_1} \right)^{-0.35}. \quad (36)$$

Therefore, the linear damping ratio modulation following damages of the wings yields to:

$$\frac{\xi_{s2}}{\xi_{s1}} \propto \left( \frac{I_2}{I_1} \right)^{0.483}. \quad (37)$$

##### S4.4 Scaling of HEA motor self-oscillation frequency with wing damage

Following the assumption that the linear damping ratio is proportional to wing inertia, in the constant motor stiffness (*i.e.*, constant or negligible muscular storage modulus) scenario, the self-oscillation frequency scaling of the HEA model with wing damage may be expressed as:

$$\frac{\Omega_{g,\text{dam}}^-}{\Omega_{g,\text{int}}^-} = \sqrt{\frac{I_{w,\text{int}}}{I_{w,\text{dam}}}} \frac{\mathcal{G}^-(\alpha, \xi_s(I_{w,\text{dam}}))}{\mathcal{G}^-(\alpha, \xi_s(I_{w,\text{int}}))}. \quad (38)$$

For this case, there is a clean bound on $\xi_p = \alpha \xi_s$ to have $\mathcal{G}^-(\alpha, \xi_s(I_w))$ real finite values. Accordingly, the exact physically relevant condition for a given $\alpha$ is:

$$\xi_p \leq \frac{\alpha}{2} \left( 1 - \frac{1}{\sqrt{\alpha^2 + 1}} \right). \quad (39)$$

The corresponding existence map in the $(\alpha, \xi_p)$ parameter space, distinguishing the real-valued and non-existent regions of $\mathcal{G}^-$, is provided as a MATLAB figure (existence\_map\_interactive.fig) in the code folder. For candidate values $\alpha \in \{1.4, 1.45, 1.5, 1.55, 2, 3\}$, the most restrictive case occurring at $\alpha = 1.4$, makes the corresponding limit that guarantees $\mathcal{G}^-$ stays real over the whole post-damaged inertia range is:

$$\xi_p \leq 0.29. \quad (40)$$

When the motor stiffness shows the maximal linear scaling associated with the muscle storage modulus (*i.e.*, muscle stiffness dominates exoskeletal stiffness), the stiffness parameter $\alpha$ behaves as frequency dependent. Then, we may express the frequency-damage scaling as:

$$\frac{\Omega_{g,dam}^-}{\Omega_{g,int}^-} = \frac{\alpha(\Omega_{int})}{\alpha(\Omega_{dam})} \sqrt{\frac{I_{w,int}}{I_{w,dam}}} \frac{\mathcal{G}^-(\alpha(\Omega_{dam}), \xi_s(I_{w,dam}))}{\mathcal{G}^-(\alpha(\Omega_{int}), \xi_s(I_{w,int}))}, \quad (41)$$

in which $\alpha$ ratio term is defined based on the muscle storage modulus trend presented in Eqn 9. in the main text:

$$\alpha_{ratio} = \frac{\alpha(\Omega_{int})}{\alpha(\Omega_{dam})} = \sqrt{\frac{k_p(\Omega_{dam})}{k_p(\Omega_{int})}} \quad (42)$$

$$= \begin{cases} \sqrt{1 + \mathcal{G}(\Omega - \Omega_{ref})} & \Omega_{max} \geq \Omega \geq \Omega_{ref} \\ \sqrt{1 + \mathcal{G}(\Omega_{max} - \Omega_{ref})} & \Omega > \Omega_{max} \end{cases},$$

Thus $\alpha_{ratio} = 1$ occurs at the intact reference state. As clipping increases resonant frequency, $\alpha_{ratio}$ decreases monotonically until $\Omega_{max}$ is reached. Beyond $\Omega_{max}$, the $\alpha_{ratio}$ remains unchanged at its minimum value. Thus, this second case incorporates both inertial reduction and stiffness modulation effects within the hybrid framework.

In this second case, to guarantee that $\mathcal{G}^-(\alpha(\Omega), \xi_s(I_w))$ stays real after damage, the condition becomes:

$$\xi_p \leq \frac{\alpha(\Omega)}{2} \left( 1 - \frac{1}{\sqrt{\alpha(\Omega)^2 + 1}} \right), \text{ for all admissible } \Omega \quad (43)$$

Therefore, the exact bound is:

$$\xi_p \leq \frac{\alpha_{min}}{2} \left( 1 - \frac{1}{\sqrt{\alpha_{min}^2 + 1}} \right), \quad \alpha_{min} = \frac{\alpha_{int}}{\max_{\Omega} \alpha_{ratio}(\Omega)}, \quad (44)$$

where $\max_{\Omega} \alpha_{ratio}(\Omega) = 1$, based on relations 42. To make it valid for all chosen intact values $\alpha_{int} \in \{1.4, 1.45, 1.5, 1.55, 2, 3\}$:

$$\xi_p \leq \min_{\alpha_{int} \in \{1.4, 1.45, 1.5, 1.55, 2, 3\}} \frac{\alpha_{int}}{2} \left( 1 - \frac{1}{\sqrt{\alpha_{int}^2 + 1}} \right). \quad (45)$$

Since this expression increases with $\alpha_{int}$, the most restrictive case is again the smallest intact value $\alpha_{int} = 1.4$. Therefore, the final exact condition for the damping ratio is equal to the previous case,

$\xi_p \leq 0.29$. The existence map provided in `existence_map_interactive.fig`, serves as a confirmation of this mathematical analysis.

To evaluate the self-oscillation frequency scaling with wing-inertia variation for a representative hybrid HEA system under the frequency-dependent muscle complex modulus scenario, Eqn 41. is solved numerically. A direct analytical solution is impractical because the parameter $\alpha$ appearing on the right-hand side of the equation is frequency dependent, while the frequency itself is the unknown quantity being solved for. The governing relation is therefore implicit, requiring the frequency and the associated stiffness modulation to be determined iteratively. To address this, a sequential piecewise numerical procedure is adopted. The damaged-to-intact wing-inertia ratio is arranged into consecutive classes, each class representing a segment of the inertia-ratio domain. Within each segment, $\alpha_{\text{ratio}}$ is assumed constant, while it is allowed to change from one segment to the next. This approximation converts the fully implicit problem into a sequence of locally explicit evaluations.

The procedure begins at the intact reference condition, where $\alpha_{\text{ratio}} = 1$. Using this value, the frequency ratios within the first inertia-ratio segment are computed from Eqn 41. The last computed frequency of that segment is then taken as the best estimate of the frequency state at the onset of the following segment. This updated frequency is substituted into Eqn 42 to determine the corresponding $\alpha_{\text{ratio}}$ for the next segment. The same marching procedure is then repeated sequentially from one segment to the next. In this way, the frequency-dependent stiffness modulation is propagated progressively across the inertia-ratio range, while preserving the causal dependence of $\alpha_{\text{ratio}}$ on the previously reached state. This sequential update is continued until the damaged frequency reaches the limiting value $\Omega_{\text{max}}$. The corresponding inertia ratio defines the threshold $I_{\text{ratio,threshold}}$: beyond this threshold, according to Eqn 42, the muscular stiffness is assumed to have reached its minimum effective level, and $\alpha_{\text{ratio}}$ therefore remains fixed at its minimum value. For all inertia ratios below $I_{\text{ratio,threshold}}$ the frequency ratio is then computed using this constant saturated value, without further segment-to-segment updates.

In the numerical implementation, the inertia-ratio domain from the intact state down to $I_{\text{ratio,threshold}}$ is partitioned into consecutive classes containing a fixed number of sampled inertia-ratio values. This grouping defines the piecewise-constant $\alpha_{\text{ratio}}$ segments used in the marching solution. The results are then displayed as staircase-like curves, where each segment corresponds to a locally constant $\alpha_{\text{ratio}}$, and the final saturated portion, where $\alpha_{\text{ratio}} = \alpha_{\text{ratio,min}}$, appears as a continuous tail. This representation makes the sequential segment-to-segment update visually explicit, while also clearly indicating the transition to the stiffness-saturated regime. Using this method, the frequency-damage scale is computed for a representative hybrid HEA system with $\xi_{p,\text{int}} = 0.29$ for five intact stiffness parameter, $\alpha_{\text{int}} \in \{1.4, 1.45, 1.5, 1.55, 2, 3\}$. The corresponding results are presented in Fig. 10 in the main text.
